# Maximum likelihood estimation of natural selection and allele age from time series data of allele frequencies

**DOI:** 10.1101/837310

**Authors:** Zhangyi He, Xiaoyang Dai, Mark Beaumont, Feng Yu

## Abstract

Temporally spaced genetic data allow for more accurate inference of population genetic parameters and hypothesis testing on the recent action of natural selection. In this work, we develop a novel likelihood-based method for jointly estimating selection coefficient and allele age from time series data of allele frequencies. Our approach is based on a hidden Markov model where the underlying process is a Wright-Fisher diffusion conditioned to survive until the time of the most recent sample. This formulation circumvents the assumption required in existing methods that the allele is created by mutation at a certain low frequency. We calculate the likelihood by numerically solving the resulting Kolmogorov backward equation backwards in time while re-weighting the solution with the emission probabilities of the observation at each sampling time point. This procedure reduces the two-dimensional numerical search for the maximum of the likelihood surface for both the selection coefficient and the allele age to a one-dimensional search over the selection coefficient only. We illustrate through extensive simulations that our method can produce accurate estimates of the selection coefficient and the allele age under both constant and non-constant demographic histories. We apply our approach to re-analyse ancient DNA data associated with horse base coat colours. We find that ignoring demographic histories or grouping raw samples can significantly bias the inference results.

## 1. Introduction

Recent advances in ancient DNA (aDNA) preparation and sequencing techniques have made available an increasing amount of high-quality time serial samples of segregating alleles in ancestral populations, *e.g.*, for humans (Sverrisdóttir et al., 2014; Mathieson et al., 2015), chickens (Flink et al., 2014; Loog et al., 2017) and horses (Ludwig et al., 2009; Pruvost et al., 2011). Such time series genetic data provide valuable information about allele frequency trajectories through time and allow for a better understanding of the evolutionary history of populations (see Leonardi et al., 2017, for a detailed review of the most recent findings in aDNA). One of the most important applications of aDNA is to study natural selection since it enables us to directly track the change in allele frequencies over time, which is the characteristic of the action of natural selection. A number of studies over the last decade have been published capitalising on the temporal aspect of aDNA data to characterise the process of natural selection. For example, Mathieson et al. (2015) used aDNA data to identify candidate loci under natural selection in European humans.

This line of work was initiated by Bollback et al. (2008), which proposed a likelihood-based approach to estimate selection coefficient from time series data of allele frequencies, assuming a Wright-Fisher model introduced by Fisher (1922) and Wright (1931). The allele frequency of the underlying population was modelled as a latent variable in a hidden Markov model (HMM), where the allele frequency of the sample drawn from the underlying population at each given time point was treated as a noisy observation of the latent population allele frequency. Since there is no tractable analytical form for the transition probabilities of the Wright-Fisher model and its numerical evaluation is computationally prohibitive for large population sizes and evolutionary timescales, the Wright-Fisher model in their likelihood calculations was approximated with its standard diffusion limit, termed the Wright-Fisher diffusion. The transition probabilities of the allele frequencies were calculated by numerically solving the Kolmogorov backward equation (KBE) resulting from the Wright-Fisher diffusion. Using the method of Bollback et al. (2008), Ludwig et al. (2009) analysed the aDNA data associated with horse coat colouration and found that natural selection acted strongly on the gene encoding the Agouti signalling peptide (*ASIP*) and the gene encoding the melanocortin 1 receptor (*MC1R*).

Malaspinas et al. (2012) extended the HMM framework of Bollback et al. (2008) to jointly estimate selection coefficient, population size and allele age based on time series data of allele frequencies. Allele age is the elapsed time since the allele was created by mutation. Along with the selection coefficient, allele age plays an important role in determining the sojourn time of a beneficial mutation (see Slatkin & Rannala, 2000, for a review). The joint estimation of allele age circumvents the assumption required in Bollback et al. (2008) that the allele frequency of the underlying population at the initial sampling time point was taken to be the observed allele frequency of the sample or was assumed to be uniformly distributed. In Malaspinas et al. (2012), the transition probabilities of the allele frequencies were calculated by approximating the Wright-Fisher diffusion with a one-step Markov process on a grid. Steinrücken et al. (2014) presented an extension of the HMM framework of Bollback et al. (2008) by capitalising on a spectral representation of the Wright-Fisher diffusion introduced by Song & Steinrücken (2012), which allows for a more general diploid model of natural selection such as the case of under- or overdominance. Ferrer-Admetlla et al. (2016) extended the HMM framework of Bollback et al. (2008) by approximating the Wright-Fisher diffusion with a coarse-grained Markov model that preserves the long-term behaviour of the Wright-Fisher diffusion. Their method can additionally estimate mutation rate. However, the methods of Steinrücken et al. (2014) and Ferrer-Admetlla et al. (2016) are unable to infer allele age as recurrent mutations were allowed in their model. Malaspinas (2016) provided an excellent review of existing approaches for studying natural selection with aDNA samples.

More recently, Schraiber et al. (2016) developed a Bayesian method under the HMM framework of Bollback et al. (2008) for the joint inference of natural selection and allele age from temporally spaced samples. Their key innovation was to apply a high-frequency path augmentation approach to circumvent the difficulty inherent in calculating the transition probabilities of the allele frequencies under the Wright-Fisher diffusion. Markov chain Monte Carlo (MCMC) techniques were employed to integrate over all possible allele frequency trajectories of the underlying population consistent with the observations. The computational advantage of the path augmentation method allows for general diploid models of natural selection and non-constant population sizes. However, to update the sample paths of the Wright-Fisher diffusion bridge, they used Bessel bridges of order four, which is somewhat challenging from both a mathematical and a programming perspective.

In the present work, we propose a novel likelihood-based approach for jointly estimating selection coefficient and allele age from time serial samples of segregating alleles. Our method is also an extension of Bollback et al. (2008) but differs from most existing approaches in two respects. Firstly, we incorporate a non-constant population size into the Wright-Fisher diffusion. Secondly, we condition the Wright-Fisher diffusion, which is taken to be the underlying process in our HMM framework, to survive until the time of the most recent sample. Our conditioned Wright-Fisher diffusion allows us to take different demographic histories into account and avoid the somewhat arbitrary initial condition that the allele was created by mutation at a certain low frequency, *e.g.*, as used in Malaspinas et al. (2012) and Schraiber et al. (2016). Our likelihood computation is carried out by numerically solving the KBE associated with the conditioned Wright-Fisher diffusion backwards in time and re-weighting the solution with the emission probabilities of the observation at each given time point. The values of the re-weighted solution at frequency 0 give the likelihood of the selection coefficient and the allele age. For each fixed selection coefficient, the likelihood for all values of the allele age can be obtained by numerically solving the KBE only once. This advance enables a reduction of the two-dimensional numerical search for the maximum of the likelihood surface for both the selection coefficient and the allele age, *e.g.*, as used in Malaspinas et al. (2012), to a one-dimensional numerical search over the selection coefficient only.

We evaluate the performance of our method with extensive simulations and show that our method allows for efficient and accurate estimation of natural selection and allele age from allele frequency time series data under both constant and non-constant demographic histories, even if the samples are sparsely distributed in time with small uneven sizes. Our simulation studies illustrate that ignoring demographic history does not affect the inference of natural selection but bias the estimation of allele age. We also use our approach to re-analyse the time serial samples of segregating alleles associated with horse base coat colours from earlier studies of Ludwig et al. (2009), Pruvost et al. (2011) and Wutke et al. (2016). We choose this aDNA dataset, despite being the focus of previous analyses (Ludwig et al., 2009; Malaspinas et al., 2012; Steinrücken et al., 2014; Schraiber et al., 2016), because it allows an instructive comparison with existing methods. Unlike these previous studies, our analysis is performed on the raw samples (drawn at 62 sampling time points) rather than the grouped samples (drawn at 9 sampling time points).

Our results suggest that horse base coat colour variation could be associated with adaptation to the climate change caused by the transition from a glacial period to an interglacial period. Finally, we perform an empirical study demonstrating that grouping aDNA samples can alter the results of the inference of natural selection and allele age.

## 2. Materials and Methods

In this section, we begin with a brief review of the Wright-Fisher diffusion for a single locus evolving subject to natural selection and then derive the Wright-Fisher diffusion conditioned to survive until a given time point. We also describe our likelihood-based method for co-estimating selection coefficient and allele age from time series data of allele frequencies, *e.g.*, how to set up the HMM framework incorporating the conditioned Wright-Fisher diffusion and how to calculate the likelihood for the population genetic quantities of interest.

### 2.1. Wright-Fisher diffusion

We consider a population of *N* randomly mating diploid individuals at a single locus *𝒜* evolving subject to natural selection according to the Wright-Fisher model (see, *e.g.*, Durrett, 2008, for more details). We assume discrete time, non-overlapping generations and non-constant population size. Suppose that there are two possible allele types at locus *𝒜*, labelled *𝒜*_1_ and *𝒜*_2_, respectively. The symbol *𝒜*_1_ is attached to the mutant allele, which is assumed to arise only once at time *t*_0_ within the population and be favoured by natural selection once it exists. The symbol *𝒜*_2_ is attached to the ancestral allele, which is assumed to originally exist in the population. Suppose that natural selection takes the form of viability selection, where viability is fixed from the time when the mutant allele arose in the population. We take relative viabilities of the three possible genotypes *𝒜*_1_*𝒜*_1_, *𝒜*_1_*𝒜*_2_ and *𝒜*_2_*𝒜*_2_ at locus *𝒜* to be 1, 1 − *hs* and 1 − *s*, respectively, where *s* ∈ [0, 1] is the selection coefficient and *h* ∈ [0, 1] is the dominance parameter.

#### 2.1.1. Wright-Fisher diffusion with selection

Let us now consider the standard diffusion limit of the Wright-Fisher model with selection. We measure time in units of 2*N*_0_ generations, denoted by *t*, where *N*_0_ is an arbitrary constant reference population size. We assume that the population size changes deterministically, with *N* (*t*) denoting the number of diploid individuals in the population at time *t*. In the diffusion limit of the Wright-Fisher model with selection, as the reference population size *N*_0_ approaches infinity, the scaled selection coefficient *α* = 2*N*_0_*s* is kept constant and the ratio of the population size to the reference population size *N* (*t*)*/N*_0_ converges to a function *ρ*(*t*). By an argument in Durrett (2008), the allele frequency trajectory through time converges to the diffusion limit of the Wright-Fisher model if we measure time in units of 2*N*_0_ generations and let the reference population size *N*_0_ go to infinity. We refer to this diffusion limit, denoted by *X*, as the Wright-Fisher diffusion with selection.

According to Durrett (2008), the Wright-Fisher diffusion *X* has the infinitesimal generator

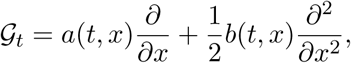

with drift term

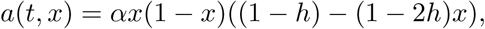

and diffusion term

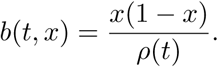

The transition probability density function of the Wright-Fisher diffusion *X*, defined by

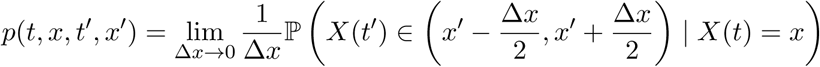

for *t*_0_ ≤ *t < t*′, can then be expressed as the solution *u*(*t, x*) of the corresponding KBE

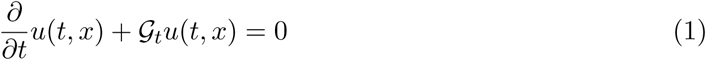

with terminal condition *u*(*t*′, ·) = *δ*(· − *x*′), where *δ* is a Dirac delta function.

#### 2.1.2. Conditioned Wright-Fisher diffusion with selection

As discussed in Valleriani (2016), it is desirable to take conditioning into account in most aDNA analyses since a single trajectory of allele frequencies through time available in aDNA results in the very limited coverage of the fitness landscape. Valleriani (2016) conditioned both the initial and the final values of the Wright-Fisher model to be fixed, assuming a finite population with perfect sampling, which is not completely realistic. In the present work, we condition the Wright-Fisher diffusion *X* to have survived until the time of the most recent sample, *i.e.*, the mutant allele frequency of the underlying population at the most recent sampling time point is strictly positive.

We let *X*^*^ denote the Wright-Fisher diffusion *X* conditioned to survive, *i.e.*, the frequency of the mutant allele stays in the interval (0, 1], until at least time *τ* after the mutant allele was created in the population at time *t*_0_. We define the transition probability density function of the conditioned Wright-Fisher diffusion *X*^*^ as

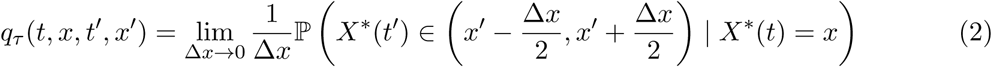

for *t*_0_ ≤ *t < t*′ *< τ*. To obtain the expression of the infinitesimal generator of *X*^*^, we need to specify the drift term, denoted by *a*^*^(*t, x*), and the diffusion term, denoted by *b*^*^(*t, x*), which are the first and second infinitesimal moments of the conditioned Wright-Fisher diffusion *X*^*^, respectively, *i.e.*,

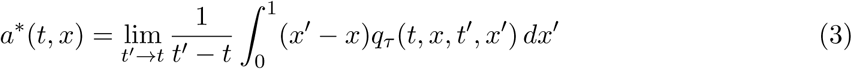

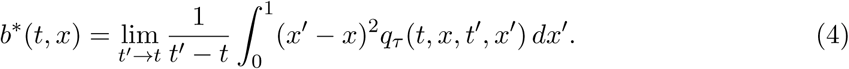

From Eq. (2), we can formulate the transition probability density function of the conditioned Wright-Fisher diffusion *X*^*^ in terms of the transition probability density function of the Wright-Fisher diffusion *X* as

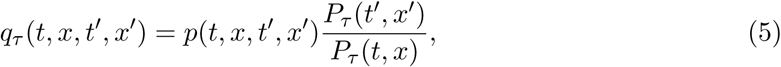

where

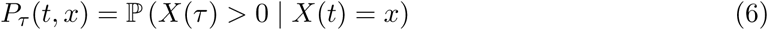

is the probability that the Wright-Fisher diffusion *X*, starting from *x* at time *t*, survives until at least time *τ*. The probability of survival, *P*_*τ*_ (*t, x*), for *t*_0_ ≤ *t < τ* can be expressed as the solution to the KBE in Eq. (1) with terminal condition 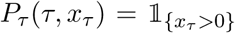, where 𝟙_*A*_ is the indicator function that equals to 1 if condition *A* holds and 0 otherwise.

Substituting Eq. (5) into Eqs. (3) and (4), we have

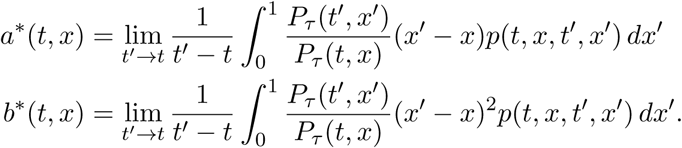

Taylor expansion yields

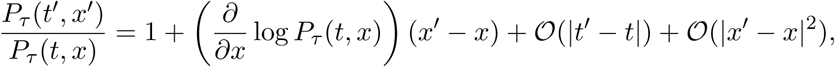

which results in

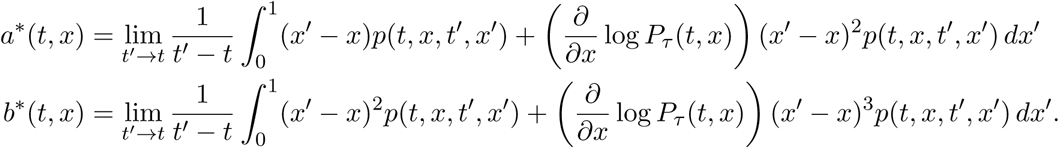

As shown in Durrett (2008), the transition probability density function of the Wright-Fisher diffusion *X* satisfies

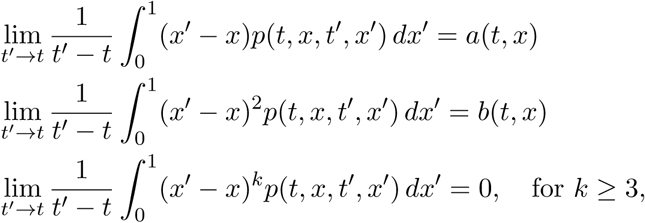

which gives rise to

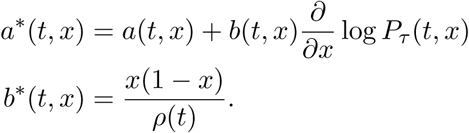

Therefore, the infinitesimal generator of the conditioned Wright-Fisher diffusion *X*^*^ can be written as

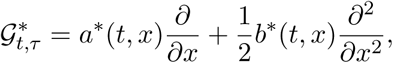

with drift term

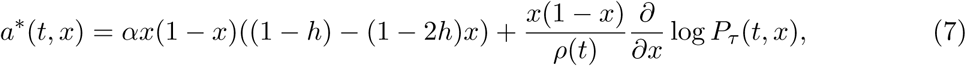

and diffusion term

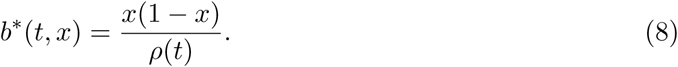

The transition probability density function of the conditioned Wright-Fisher diffusion *X*^*^ can then be expressed as the solution *u*(*t, x*) of the corresponding KBE

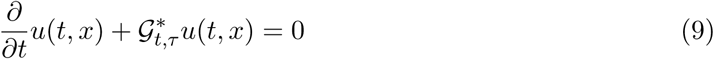

with terminal condition *u*(*t*′, ·) = *δ*(· − *x*′).

### 2.2 Maximum likelihood estimation

Suppose that the available data are always sampled from the underlying population at a finite number of distinct time points, say *t*_1_ *< t*_2_ *< … < t*_*K*_, where the time is measured in units of 2*N*_0_ generations to be consistent with the time scale of the Wright-Fisher diffusion. At the sampling time point *t*_*k*_, let *c*_*k*_ represent the number of mutant alleles observed in a sample of *n*_*k*_ chromosomes drawn from the underlying population. In this work, the population genetic quantities of interest are the scaled selection coefficient *α*, the dominance parameter *h* and the allele age *t*_0_.

#### 2.2.1. Hidden Markov model

To our knowledge, Malaspinas et al. (2012) and Schraiber et al. (2016) are the only existing works that seek to jointly infer natural selection and allele age from time series data of allele frequencies. These two approaches were both based on the HMM framework of Bollback et al. (2008) incorporating the Wright-Fisher diffusion with selection. To jointly estimate allele age, initial conditions for the Wright-Fisher diffusion *X* at time *t*_0_ must be specified. Schraiber et al. (2016) took the mutant allele frequency *X*(*t*_0_) to be some small but arbitrary value, which was found to be feasible in their approach but is slightly unsatisfying. Malaspinas et al. (2012) took the mutant allele frequency *X*(*t*_0_) to be 1*/*(2*N* (*t*_0_)), which corresponds to the case that the positively selected allele was created as a *de novo* mutation. This can be slightly problematic in that the Wright-Fisher diffusion *X* may hit frequency 1*/*(2*N* (*t*_0_)) again after the mutation arose, so the time *t* when *X*(*t*) = 1*/*(2*N* (*t*_0_)) may not be the same as the allele age.

Similar to Bollback et al. (2008), our approach is also built on an HMM framework except that the underlying population evolves according to the conditioned Wright-Fisher diffusion *X*^*^ rather than the Wright-Fisher diffusion *X*. Given the mutant allele frequency trajectory of the underlying population, the observations are modelled as independent binomial samples drawn from the underlying population at every sampling time point (see Figure 1 for the graphical representation of our HMM framework). The conditioning enables us to attach an initial condition *X*^*^(*t*_0_) = 0 with 0 forming an entrance boundary (see Supplemental Material, File S1), *i.e.*, the conditioned Wright-Fisher diffusion *X*^*^ will reach the interval (0, 1] starting from the initial mutant allele frequency *X*^*^(*t*_0_) = 0. Our setup allows us to avoid specifying any arbitrary starting mutant allele frequency of the underlying population at time *t*_0_.

**Figure 1:**
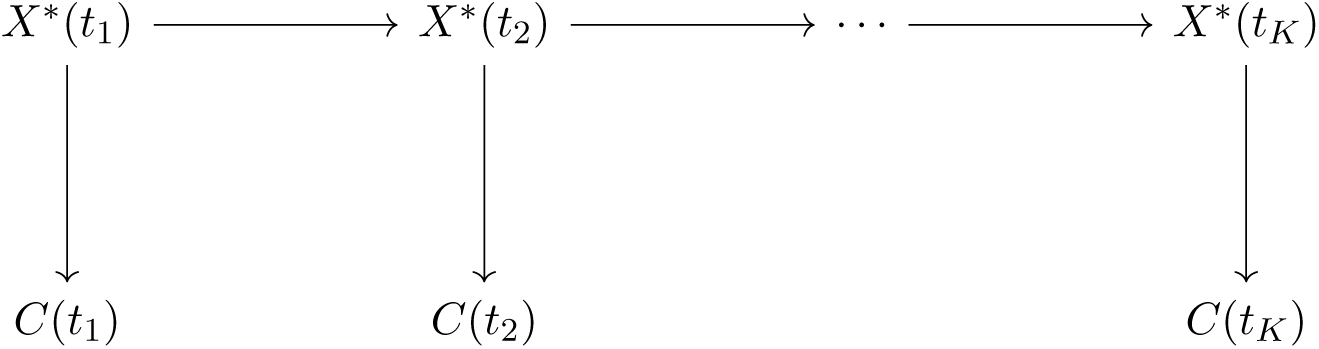
Graphical representation of the HMM framework for time series data of allele frequencies.

More specifically, our HMM framework can be fully captured by a bivariate Markov process {(*X*^*^(*t*), *C*(*t*)) : *t* ∈ [*t*_1_, *t*_*K*_]} with initial condition *X*^*^(*t*_0_) = 0, where the unobserved process *X*^*^(*t*) for *t* ∈ [*t*_1_, *t*_*K*_] is the Wright-Fisher diffusion conditioned to survive until the most recent sampling time point *t*_*K*_, and the observed process *C*(*t*) for *t* ∈ {*t*_1_, *t*_2_, *…, t*_*K*_} is a sequence of conditionally independent binomial random variables given the unobserved process *X*^*^(*t*) at each sampling time point. The transition probabilities for our HMM between two consecutive sampling time points *t*_*k*−1_ and *t*_*k*_ are

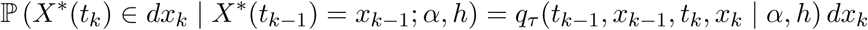

for *k* = 2, 3, *…, K*, where *q*_*τ*_ (*t*_*k*−1_, *x*_*k*−1_, *t*_*k*_, *x*_*k*_ | *α, h*) is the transition probability density function of the conditioned Wright-Fisher diffusion *X*^*^ that satisfies the KBE in Eq. (9) with terminal condition *q*_*τ*_ (*t*_*k*_, ·) = *δ*(· − *x*_*k*_). The emission probabilities for our HMM at the sampling time point *t*_*k*_ are

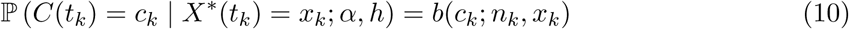

for k = 1, 2, …, K, where

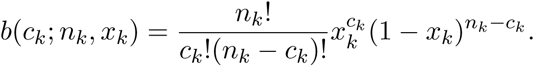

#### 2.2.2. Likelihood computation

To calculate the likelihood for the population genetic parameters of interest, defined by

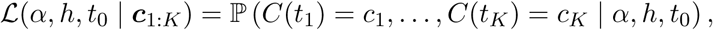

we let *𝒯*_0_ = (−∞, *t*_1_) and *𝒯*_*k*−1_ = [*t*_*k*−1_, *t*_*k*_) for *k* = 2, 3, *…, K*, and decompose the likelihood to a sum of terms according to which time interval *𝒯*_*k*−1_ the allele age *t*_0_ falls in,

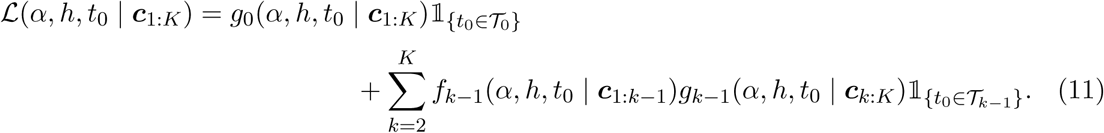

Note that in the decomposition of Eq. (11), only one term will be nonzero since the allele age *t*_0_ can only be in one of the time intervals *𝒯*_*k*−1_ for *k* = 1, 2, *…, K*. In Eq. (11), if the allele age *t*_0_ ∈ *𝒯*_*k*−1_, then

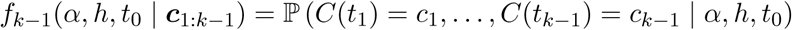

is the probability of the observations sampled before the time *t*_0_ that the mutant allele arose in the underlying population, and

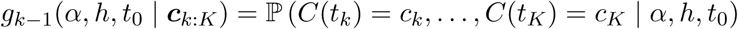

is the probability of the observations sampled after the time *t*_0_ that the mutant allele was created in the underlying population. Given that the conditioned Wright-Fisher diffusion *X*^*^(*t*) = 0 for *t* ∈ (−∞, *t*_0_], we have

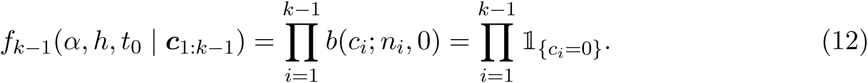

Under our HMM framework, we have

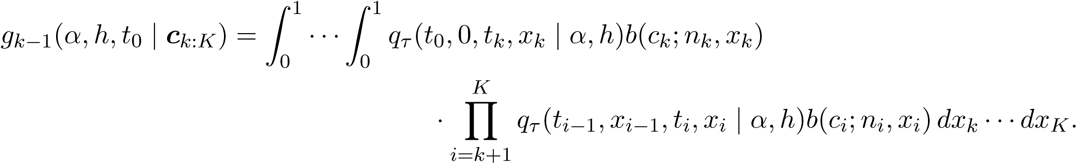

We define

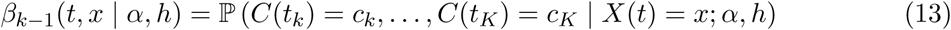

to be the probability of the observations sampled after the time *t* ∈ *𝒯* _*k*−1_ given initial condition *X*^*^(*t*) = *x*, which can be calculated recursively by

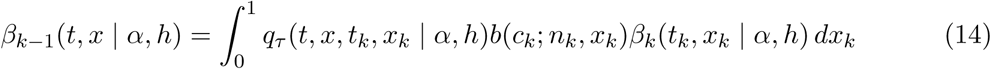

for *k* = *K,K* − 1, …, 1, given terminal condition

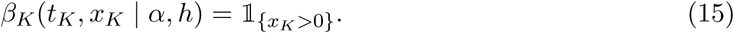

The recursive formula in Eq. (14) implies that the probability *β*_*k*−1_(*t, x* | *α, h*) for *t* ∈ *𝒯*_*k*−1_ can be expressed as the solution to the KBE in Eq. (9) with terminal condition

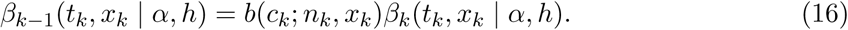

From Eq. (13), we have

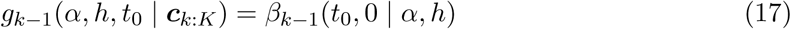

for *k* = 1, 2, *…, K*. Substituting Eqs. (12) and (17) into Eq. (11), we can formulate the likelihood for the population genetic parameters of interest as

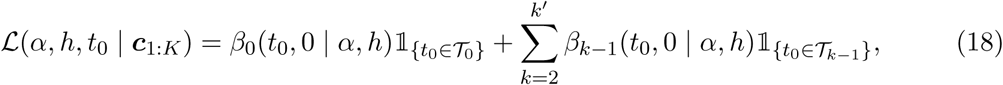

where

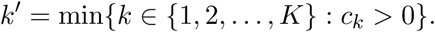

From the calculation leading to Eq. (18), the likelihood for all possible values of the allele age *t*_0_ with fixed values of the scaled selection coefficient *α* and the dominance parameter *h* can be obtained by recursively calculating the probabilities *β*_*k*−1_(*t, x* | *α, h*) for *t* ∈ *𝒯*_*k*−1_ and *x* ∈ [0, 1] for *k* = *K, K* − 1, *…*, 1 with Eq. (14). However, the KBE in Eq. (9) for the conditioned Wright-Fisher diffusion *X*^*^ cannot be solved analytically. We resort to a finite difference approach like the Crank-Nicolson scheme proposed by Crank & Nicolson (1947) to get the numerical solution of the KBE in Eq. (9), which requires us to compute the survival probability *P*_*τ*_ (*t, x*) in Eq. (6) for *t* ∈ (−∞, *t*_*K*_] and *x* ∈ [0, 1]. For this, we numerically solve the KBE in Eq. (1) for the Wright-Fisher diffusion *X*. We then use the values of the survival probability *P*_*τ*_ (*t, x*) to numerically solve the KBE in Eq. (9). Note that we take the drift term 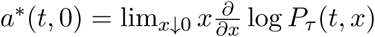 to be 1, which is justified in Supplemental Material, File S1.

For clarity, we write down the procedure we follow to obtain the likelihood *ℒ*(*α, h, t*_0_ | ***c***_1:*K*_) for all possible values of the allele age *t*_0_ ∈ (−∞, *t*_*K*_] with fixed values of the scaled selection coefficient *α* and the dominance parameter *h*:

Step 1: Calculate *P*_*τ*_ (*t, x*) for *t* ∈ (−∞, *t*_*K*_) and *x* ∈ [0, 1] by numerically solving the KBE in Eq. (1) backwards in time with terminal condition 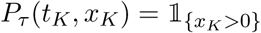 for *x*_*K*_ ∈ [0, 1] and boundary conditions *P*_*τ*_ (*t*, 0) = 0 and *P*_*τ*_ (*t*, 1) = 1 for *t* ∈ (−∞, *t*_*K*_).

Step 2: Calculate 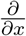 log *P*_*τ*_ (*t, x*) for *t* ∈ (−∞, *t*_*K*_) and *x* ∈ [0, 1] by numerically differentiating log *P*_*τ*_ (*t, x*).

Step 3: Initialise *β*_*K*_ (*t, x* | *α, h*) with Eq. (15).

Step 4: Set *k* = *K* and repeat until *k* = 2:

Step 4a: Update the terminal condition *β*_*k*−1_(*t*_*k*_, *x*_*k*_ | *α, h*) with Eq. (16).

Step 4b: Calculate *β*_*k*−1_(*t, x* | *α, h*) for *t* ∈ *𝒯*_*k*−1_ and *x* ∈ [0, 1] by numerically solving the KBE in Eq. (9) backwards in time with terminal condition *β*_*k*−1_(*t*_*k*_, *x*_*k*_ | *α, h*) for *x*_*k*_ ∈ [0, 1].

Step 4c: Set *k* = *k* − 1.

Step 5: Update the terminal condition *β*_0_(*t*_1_, *x*_1_ | *α, h*) with Eq. (16).

Step 6: Calculate *β*_0_(*t, x* | *α, h*) for *t* ∈ *𝒯*_0_ and *x* ∈ [0, 1] by numerically solving the KBE in Eq. (9) backwards in time with terminal condition *β*_0_(*t*_1_, *x*_1_ | *α, h*) for *x*_1_ ∈ [0, 1] until 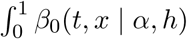 is sufficiently small.

Step 7: Combine *β*_*k*−1_(*t*_0_, 0 | *α, h*) for *k* = 1, 2, *…, k*′ using Eq. (18) to obtain *ℒ*(*α, h, t*_0_ | ***c***_1:*K*_).

To obtain the maximum of the likelihood for the population genetic quantities of interest, we perform a section search over the scaled selection coefficient and the dominance parameter. We start with a fixed region for all possible values of the scaled selection coefficient and the dominance parameter. For each pair of the *m* equally spaced scaled selection coefficients, *α*_1_, *α*_2_, *…, α*_*m*_, and the *n* equally spaced dominance parameters, *h*_1_, *h*_2_, *…, h*_*n*_, within the region, we perform the procedure laid out above to obtain the likelihood for all possible values of the allele age. We record the maximum value of the likelihood as well as the allele age where this maximum is attained. Suppose that the scaled selection coefficient *α*_*i*′_ and the dominance parameter *h*_*j*′_ yield the highest likelihood amongst the combinations of the scaled selection coefficient *α*_*i*_ for *i* = 1, 2, *… m* and the dominance parameter *h*_*j*_ for *j* = 1, 2, *… n*. If the highest likelihood is achieved at an interior point, in the next step, we narrow our search region to [*α*_*i′*−1_, *α*_*i′*+1_] *×* [*h*_*j′*−1_, *h*_*j′*+1_]. Otherwise, *i.e.*, if the highest likelihood is achieved at a boundary point, we shift the search grid to centre at the parameters values where the likelihood achieves its maximum from the previous step. We continue this procedure until the area of the search region is sufficiently small.

### 2.3. Data availability

Supplemental material available at FigShare. Source code for the method described in this work is available at https://github.com/zhangyi-he/WFM-1L-DiffusApprox-KBE.

## 3. Results

In this section, we first evaluate the performance of our method through a number of simulated datasets with given population genetic parameter values. We then employ our approach to analyse aDNA time series data of allele frequencies associated with horse coat colouration from previous studies of Ludwig et al. (2009), Pruvost et al. (2011) and Wutke et al. (2016), where eight coat colour genes from ancient horse samples ranging from a pre- to a post-domestication period were sequenced.

### 3.1. Analysis of simulated data

To assess the performance of our approach, we simulate sets of data with different population genetic parameter values. For each simulated dataset, we specify the demographic history *N* (*k*) and fix the selection coefficient *s*, the dominance parameter *h* and the allele age *k*_0_, where *k* and *k*_0_ are measured in generations. We simulate mutant allele frequency trajectories of the underlying population using the Wright-Fisher model with selection (see, *e.g.*, Durrett, 2008) with initial frequency 1*/*(2*N* (*k*_0_)). Given the realisation of the mutant allele frequency trajectory of the underlying population, we generate the mutant allele count of the sample independently at each sampling time point according to the binomial distribution in Eq. (10). We only keep simulated datasets where the mutant allele survives in the underlying population until the time of the most recent sample.

As in Malaspinas et al. (2012), we consider two different sampling schemes in our empirical studies to quantify whether it is better to sample more chromosomes at fewer time points or the opposite within a given sampling period. In sampling scheme A, we draw 20 chromosomes at *K* = 60 evenly spaced sampling time points from the underlying population, whereas in sampling scheme B, we draw 120 chromosomes at *K* = 10 evenly spaced sampling time points. For each sampling scheme, we choose four basic demographic histories: the constant model, the bottleneck model, the growth model and the decline model, and fix the selection coefficient and the dominance parameter to several potential values: *s* ∈ {0, 0.0025, 0.005, 0.01, 0.015, 0.02} and *h* ∈ {0, 0.5, 1}.

Due to the vastly different values of selection coefficient and dominance parameter, it is not possible to use a single set of fixed sampling times for all simulated datasets. Instead, for each pair of given values of the selection coefficient and the dominance parameter, we perform the following steps to determine the sampling time points:

Step 1: Generate 100 realisations of the mutant allele frequency trajectory of the underlying population through the Wright-Fisher model with selection.

Step 2: Take the mean of the 100 realisations of the mutant allele frequency trajectory of the underlying population.

Step 3: Set the time when the mean mutant allele frequency trajectory of the underlying population first hits frequency 0.1 and 0.9 to be the first and last sampling time points, labelled *k*_1_ and *k*_*K*_, respectively.

Step 4: Set the rest of the sampling time points *k*_2_, *k*_3_ *…, k*_*K*−1_ to be evenly spaced between the first and last sampling time points *k*_1_ and *k*_*K*_.

Under this scheme, a significant number of the realisations of the mutant allele frequency trajectory of the underlying population will capture a significant proportion of the selective sweep. See Figures 2 and 3 for the simulated datasets under constant and bottleneck demographic histories for each sampling scheme with their corresponding likelihood surfaces produced through our approach as representative examples. Note that in what follows we set the reference population size *N*_0_ = 16000 unless otherwise noted

**Figure 2:**
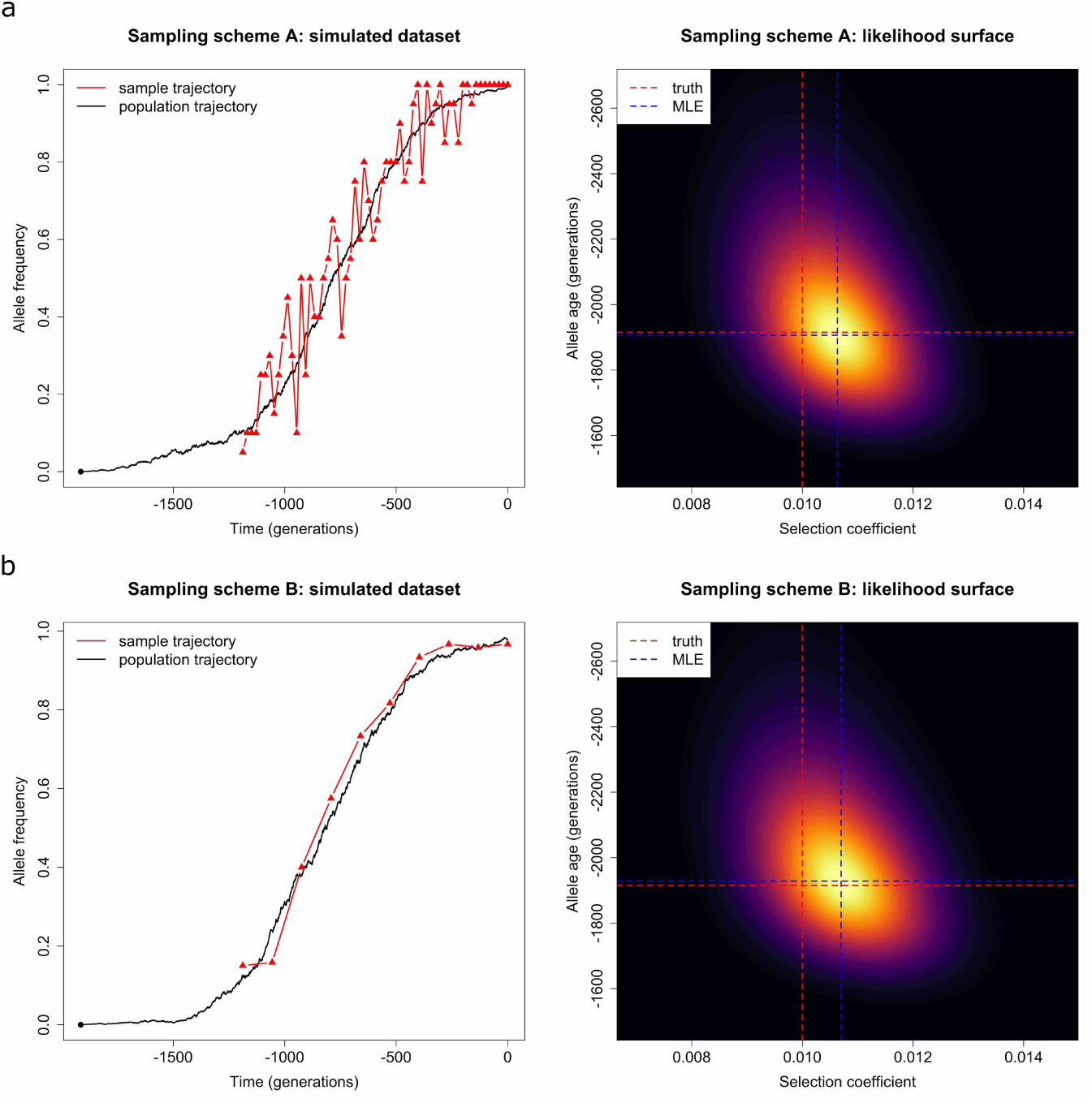
Representative examples of the datasets simulated under a constant demographic history for every sampling scheme with their resulting likelihood surfaces. We take the selection coefficient to be *s* = 0.01 and the dominance parameter to be *h* = 0.5. We set the population size to *N* (*k*) = 16000 for *k* ∈ {−1915, −1914, *…*, 0}. The mutant allele is created at frequency *x*_0_ = 3.125 *×* 10^−5^ in the underlying population at time *k*_0_ = −1915 (black filled circle), and the mutant allele frequency trajectory of the underlying population is simulated through the Wright-Fisher model with selection (black line). (a) Sampling scheme A: 20 chromosomes drawn from the underlying population every 20 generations (red filled triangle) from generation -1188 to 0. (b) Sampling scheme B: 120 chromosomes drawn from the underlying population every 132 generations (red filled triangle) from generation -1188 to 0.

**Figure 3:**
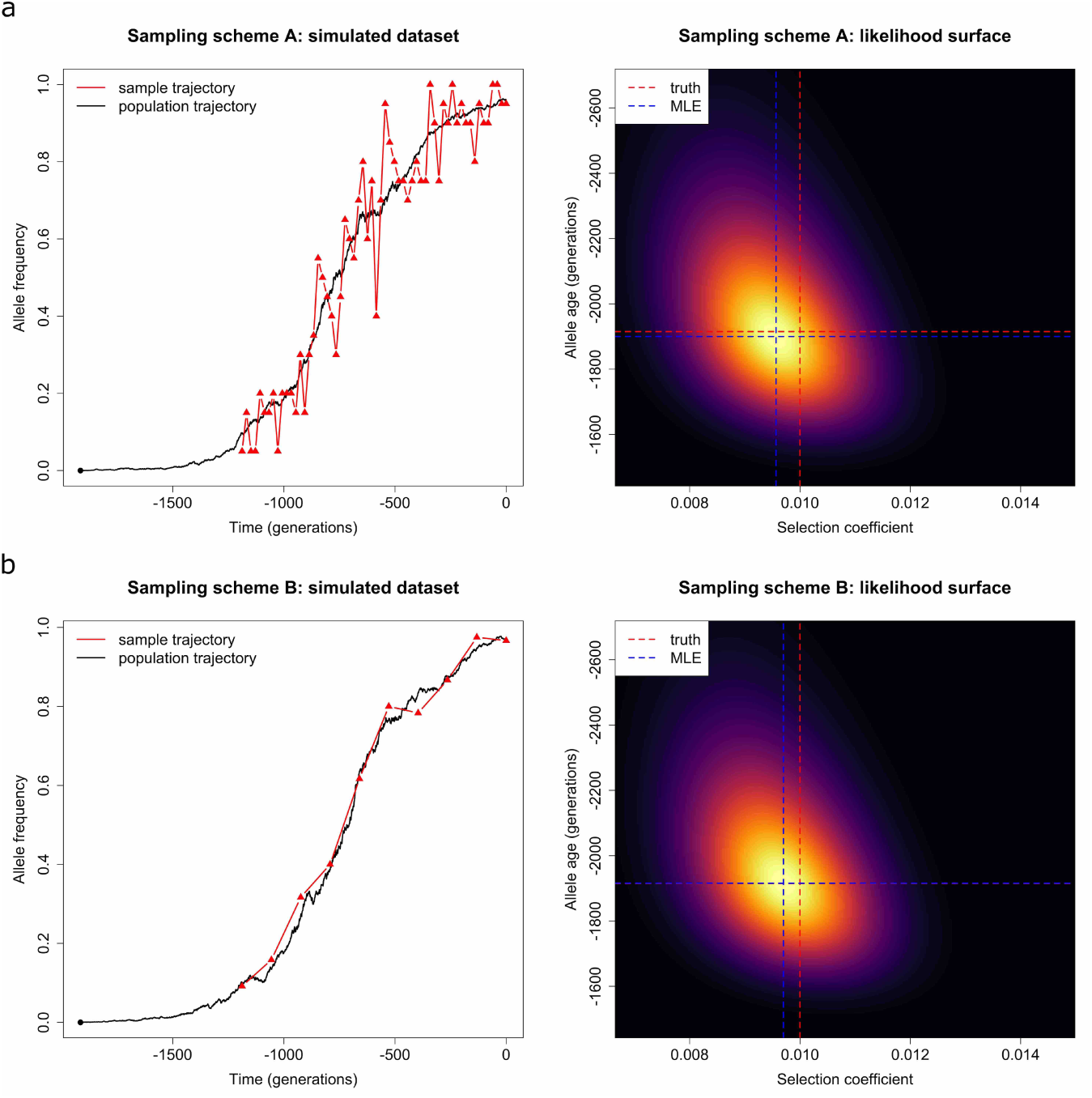
Representative examples of the datasets simulated under a bottleneck demographic history for every sampling scheme with their resulting likelihood surfaces. We simulate the mutant allele frequency trajectory of the underlying population according to the Wright-Fisher model with selection, where the population size is taken to be *N* (*k*) = 8000 for *k* ∈ {−1436, −1435, *…*, −957} and *N* (*k*) = 16000 otherwise, and the rest of the parameters are the same as those in Figure 2.

#### 3.1.1. Power to infer natural selection and allele age

We first test the accuracy of our approach across different parameter values and sampling schemes under a constant demographic history, where *N* (*k*) = 16000 for *k* ∈ 𝕫. We present the boxplot results for the resulting estimates for these simulated datasets across different parameter values and sampling schemes in Figure 4. The tips of the whiskers represent the 2.5%-quantile and the 97.5%-quantile, and the boxes denote the first and third quartiles with the median in the middle. We summarise the bias and the root mean square error (RMSE) of the resulting estimates in Supplemental Material, Tables S1-S3.

**Figure 4:**
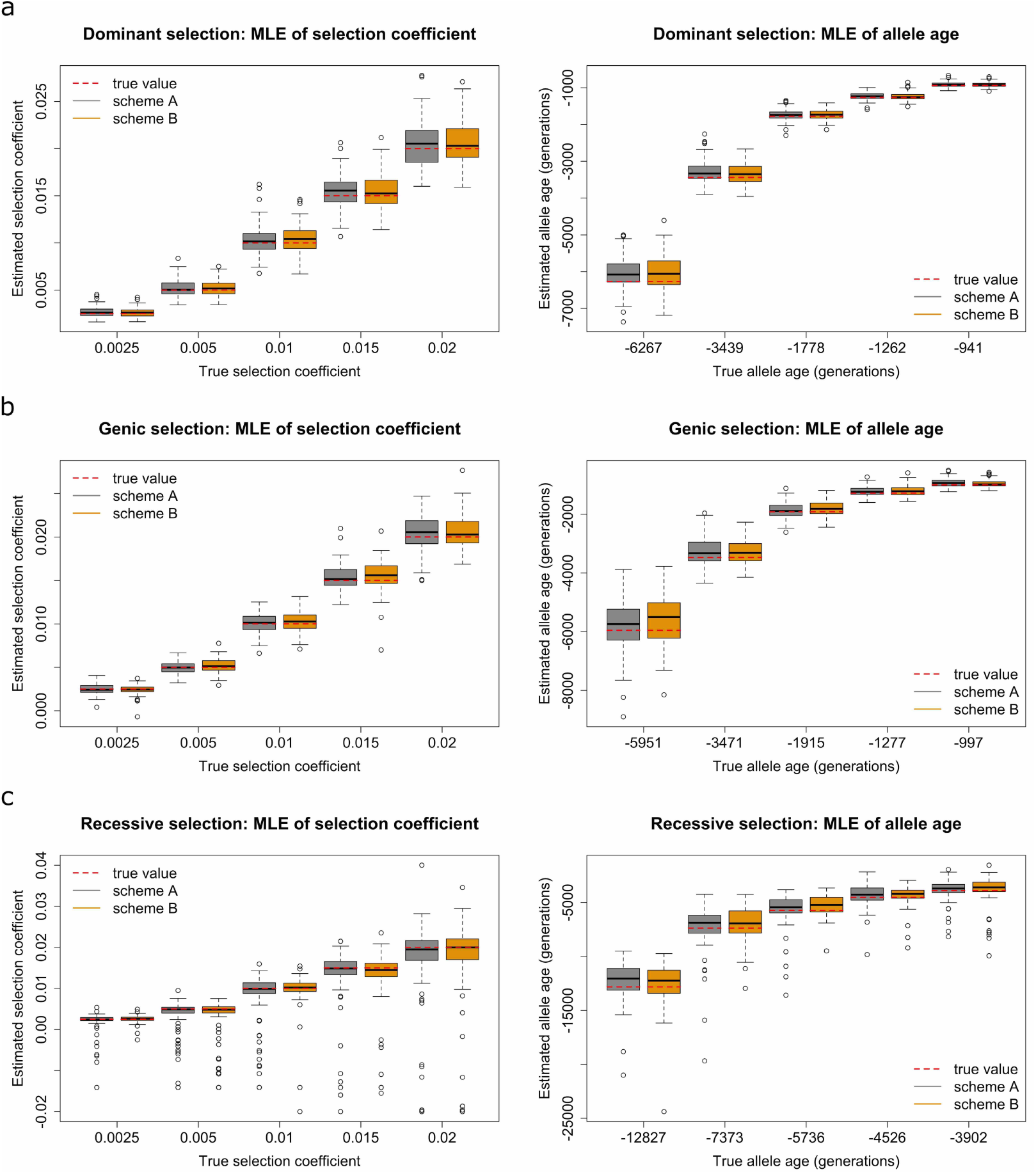
Empirical distributions of estimates for 100 datasets simulated under a constant demographic history with population size *N* (*k*) = 16000 for *k* ∈ {*k*_0_, *k*_0_ + 1, *…*, 0}. Grey boxplots represent the estimates produced for sampling scheme A, and yellow boxplots represent the estimates produced for sampling scheme B. Boxplots of the estimates for (a) dominant selection (*h* = 0) (b) genic selection (*h* = 0.5) (c) recessive selection (*h* = 1).

As can be observed from Figure 4, our estimates for the selection coefficient and the allele age both show little bias across different parameter values and sampling schemes, although one can discern a slight positive bias for the estimates of the allele age. Given that there is no requirement for the maximum likelihood estimator to be unbiased in general, some degree of bias is not unexpected. An additional cause of the bias arises from the finite grid size (*i.e.*, the state space [0, 1] is divided into 270 subintervals) used when we numerically solve the KBE in Eq. (9) through the Crank-Nicolson approach. Numerically solving the KBE with a finer grid results in a smaller bias, although the numerical solution takes much longer to run. With the increase of the selection coefficient, the uncertainty of the estimate for the selection coefficient increases whereas the uncertainty of the estimate for the allele age decreases. This can be explained by the relatively smaller noise with larger selection coefficients, yielding more information about the mutant allele frequency trajectory of the underlying population prior to the first sample time point. Moreover, the 2.5%-quantiles of the empirical distributions of the estimates for the selection coefficient *s >* 0 are all larger than the 97.5%-quantile of the empirical distribution of the estimates for the selection coefficient *s* = 0 (*i.e., selective neutrality*, see Supplemental Material, Figure S1), which implies that our method has a strong power to reject neutrality.

Comparing boxplot results for different selection schemes, we find that there are many more outliers found in the case of recessive selection (*h* = 1, Figure 4c) than in the cases of dominant and genic selection (*h* = 0 and *h* = 0.5, Figures 4a and 4b). To understand this effect, we plot in Figure 5 realisations of mutant allele frequency trajectories of the underlying population for all simulated datasets. To capture information on selection coefficient and allele age, the underlying mutant allele frequency trajectory should ideally grow from a lower frequency around 0 to a high frequency around 1 within the sampling period. However, as can be seen in Figure 5, there are a number of simulated mutant allele frequency trajectories of the underlying population where the mutant allele frequencies at all sampling time points are all close to the absorbing boundaries, 0 or 1. This effect is especially pronounced for the case of recessive selection (*h* = 1). Such mutant allele frequency trajectories of the underlying population contain little information about the underlying selection coefficient and allele age, therefore more outliers when we try to estimate these parameters.

**Figure 5:**
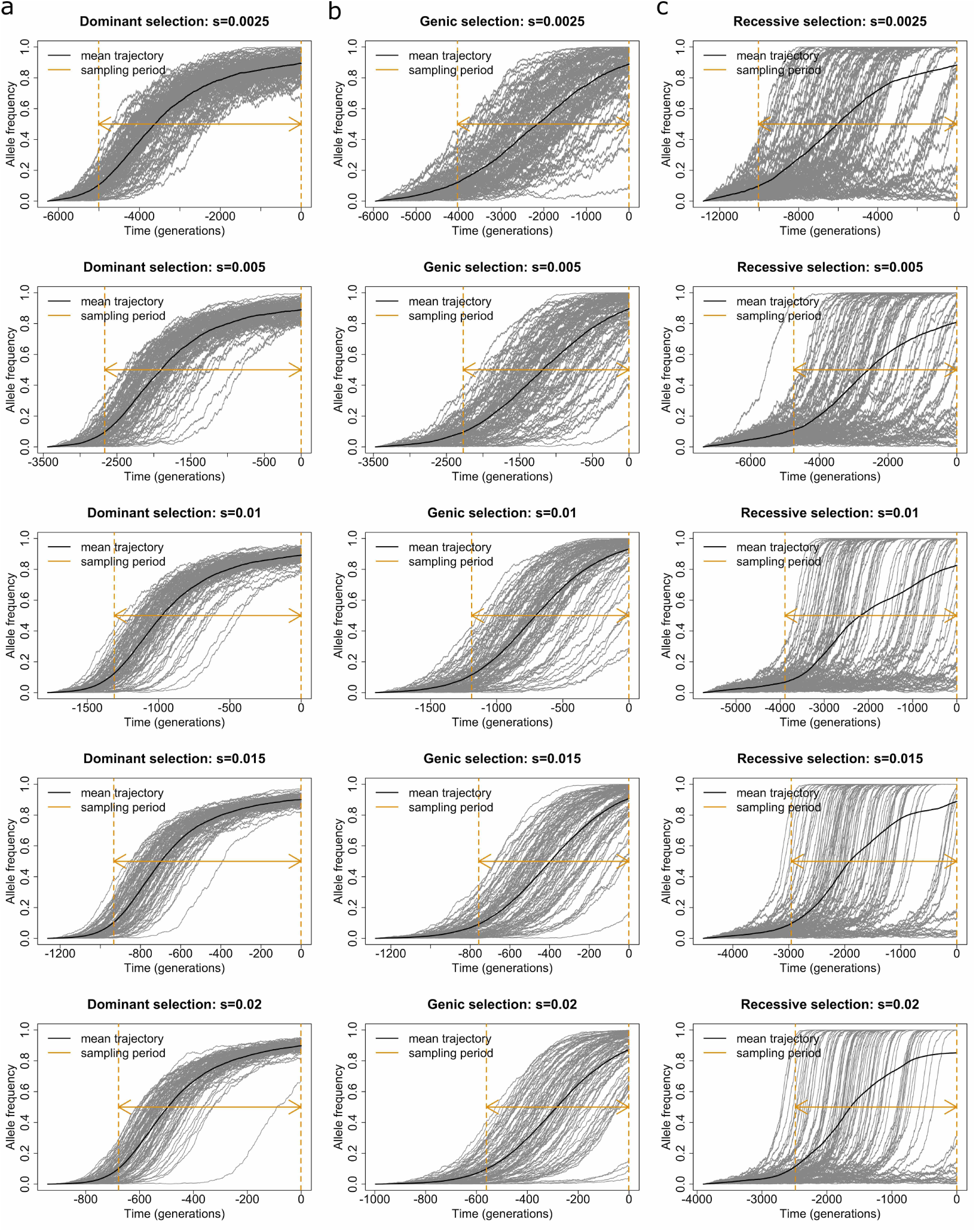
Mutant allele frequency trajectories of the underlying population for the datasets simulated under a constant demographic history with population size *N* (*k*) = 16000 for *k* ∈ {*k*_0_, *k*_0_ + 1, *…*, 0}, and dominant parameter (a) *h* = 0 (dominant selection) (b) *h* = 0.5 (genic selection) (c) *h* = 1 (recessive selection).

Comparing boxplot results under different sampling schemes, we can observe similar results for estimates of the selection coefficient and the allele age, which suggests that within a given sampling period, sampling more chromosomes at fewer time points or the opposite has little effect on the inference of natural selection and allele age. We also present the empirical distributions of the estimates for the selection coefficient and the allele age obtained from the samples sparsely distributed in time with small uneven sizes in Supplemental Material Figure S2, with their bias and RMSE summarised in Supplemental Material Table S4. These results further establish the ability of our method to handle data generated under any sampling schemes even if samples are sparsely distributed in time with small uneven sizes.

We now assess the performance of our method across different parameter values and sampling schemes under non-constant demographic histories and compare the results of the inference with and without taking demographic history into account. The resulting boxplots of the empirical studies under a bottleneck demographic history are illustrated in Figure 6 (the grey boxes), where the population size is taken to be *N* (*k*) = 8000 for *k* ∈ {[*k*_0_*/*2], [*k*_0_*/*2] + 1, *…*, [3*k*_0_*/*4]} and *N* (*k*) = 16000 otherwise. The bias and RMSE of the resulting estimates are summarised in Supplemental Material, Table S5. In order to investigate the effect of the demographic history on the inference results, we also present the boxplot results produced without incorporating the true demographic history in Figure 6 (the yellow boxes), where we fix the population size to be the bottleneck population size *N* (*k*) = 8000 for *k* ∈ {*k*_0_, *k*_0_ + 1, *…*, 0}, with all other settings being identical to the empirical studies for the bottleneck demographic history. The bias and the RMSE of the resulting estimates are summarised in Supplemental Material, Table S6.

**Figure 6:**
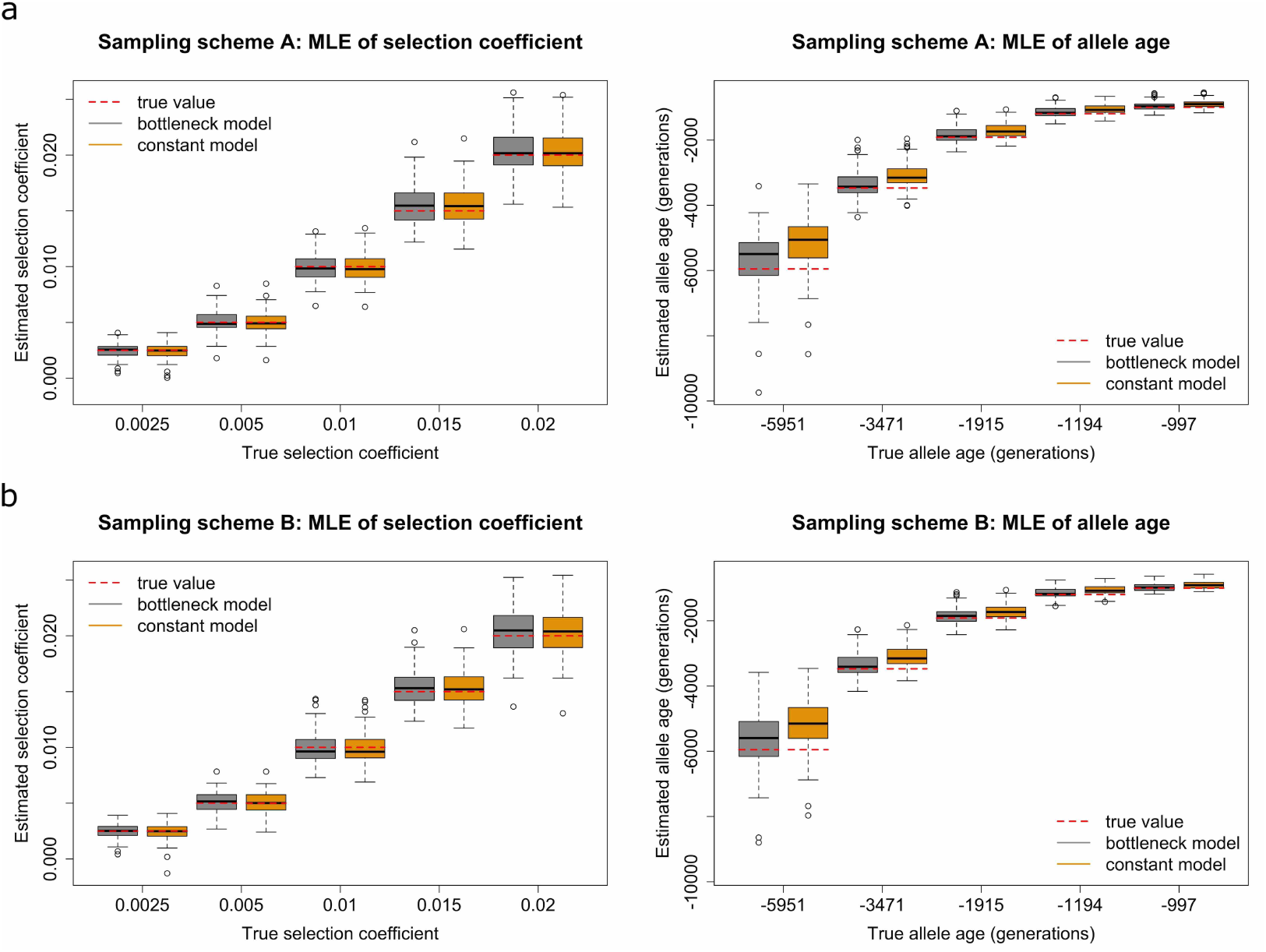
Empirical distributions of estimates for 100 datasets simulated under a bottleneck demographic history with population size *N* (*k*) = 8000 for *k* ∈ {[*k*_0_*/*2], [*k*_0_*/*2] + 1, *…*, [3*k*_0_*/*4]} and *N* (*k*) = 16000 otherwise, and dominance parameter *h* = 0.5. Grey boxplots represent the estimates assuming the true demographic history, and yellow boxplots assuming a constant demographic history with the bottleneck population size *N* (*k*) = 8000 for *k* ∈ {*k*_0_, *k*_0_ + 1, *…*, 0}. Boxplots of the estimates for (a) sampling scheme A (b) sampling scheme B.

As illustrated in Figure 6, we find that our estimates for the selection coefficient and the allele age under the bottleneck demographic history are both reasonably accurate across different parameter values and sampling schemes when we incorporate the true demographic history. They are very similar to the estimates in the empirical studies under the constant demographic history shown in Figure 4. Comparing the boxplot results produced with and without considering demographic histories, we observe that incorporating true demographic histories has little effect on the estimation of natural selection. This is consistent with the findings of Jewett et al. (2016) that accurate estimates of the selection coefficient could often be obtained by assuming that alleles evolve without genetic drift in a population of infinite size.

As can be seen in Figure 6, however, ignoring true demographic histories can cause significant biases in estimating the allele age. We will give an intuitive explanation for why this bias occurs. In this simulation study, the earliest true population size is 16000, but the inference results ignoring demographic history takes the population size to be always 8000, half as large as the true value for early times. If we take the reference population size to be *N*_0_ = 16000 under both scenarios, the drift term *a*^*^(*t, x*) in Eq. (7) under the assumption of the constant population size of 8000 will have *ρ*(*t*) = 1*/*2 for early times, larger than the drift term *a*^*^(*t, x*) if *ρ*(*t*) = 1 under the true population size of 16000. This extra upward drift causes *X*^*^, the conditioned frequency of the mutant allele, to increase more rapidly than what actually happens under true demographic history. Therefore, the resulting estimates of the allele age are smaller than the true allele age. A similar conclusion can be reached from other non-constant demographic histories. See the empirical distributions of the estimates under growth and decline demographic histories in Supplemental Material Figures S3 and S4, with their bias and RMSE summarised in Supplemental Material, Tables S7-S10, where we take the population size to be *N* (*k*) = 16000 for *k* ∈ {*k*_0_, *k*_0_ + 1, *…*, [*k*_0_*/*2]} and *N* (*k*) = 32000 otherwise for the growth demographic history and the population size to be *N* (*k*) = 100000 for *k* ∈ {*k*_0_, *k*_0_ + 1, *…*, [*k*_0_*/*2]} and *N* (*k*) = 16000 otherwise for the decline demographic history. The boxplots of the resulting estimates for the case of selective neutrality (*s* = 0) under non-constant demographic histories can be found in Supplemental Material, Figure S5.

In summary, our approach can deliver accurate joint estimates of the selection coefficient and the allele age from time series data of allele frequencies across different parameter values, demographic histories and sampling schemes, even if the samples are sparsely distributed in time with small uneven sizes. Our empirical studies show that ignoring true demographic histories has minimal impact on the inference of natural selection but significantly alters the estimation of allele age. We also find that within any given sampling period, drawing more chromosomes at fewer time points or the opposite has little effect on the estimation of the selection coefficient and the allele age. In all these empirical studies above, the dominance parameter is treated as known. We also present the results where the dominance parameter is estimated along with the selection coefficient and the allele age (see Supplemental Material, Table S11).

#### 3.1.2. Comparison with existing methods

We compare the performance of our method with the approach of Schraiber et al. (2016). To our knowledge, Schraiber et al. (2016) is the only existing method that allows for the joint inference of natural selection and allele age from time series data of allele frequencies while different demographic histories can be explicitly incorporated. We apply their approach to re-analyse the datasets generated under different demographic histories in Section 3.1.1. Following Schraiber et al. (2016), we run their MCMC method with 1000000 iterations for each simulated dataset, sampling every 1000 iterations to get 1000 MCMC samples and then discard the first 500 samples as burn-in period. We show the results of both approaches under the constant demographic history in Figure 7 and leave the results under non-demographic histories in Supplemental Material, Figures S6-S8. The bias and RMSE of the estimates are summarised in Supplemental Material, Tables S12-S15. The empirical distributions of estimates in the case of the selection coefficient *s* = 0 under different demographic histories are illustrated in Supplemental Material, Figure S9.

**Figure 7:**
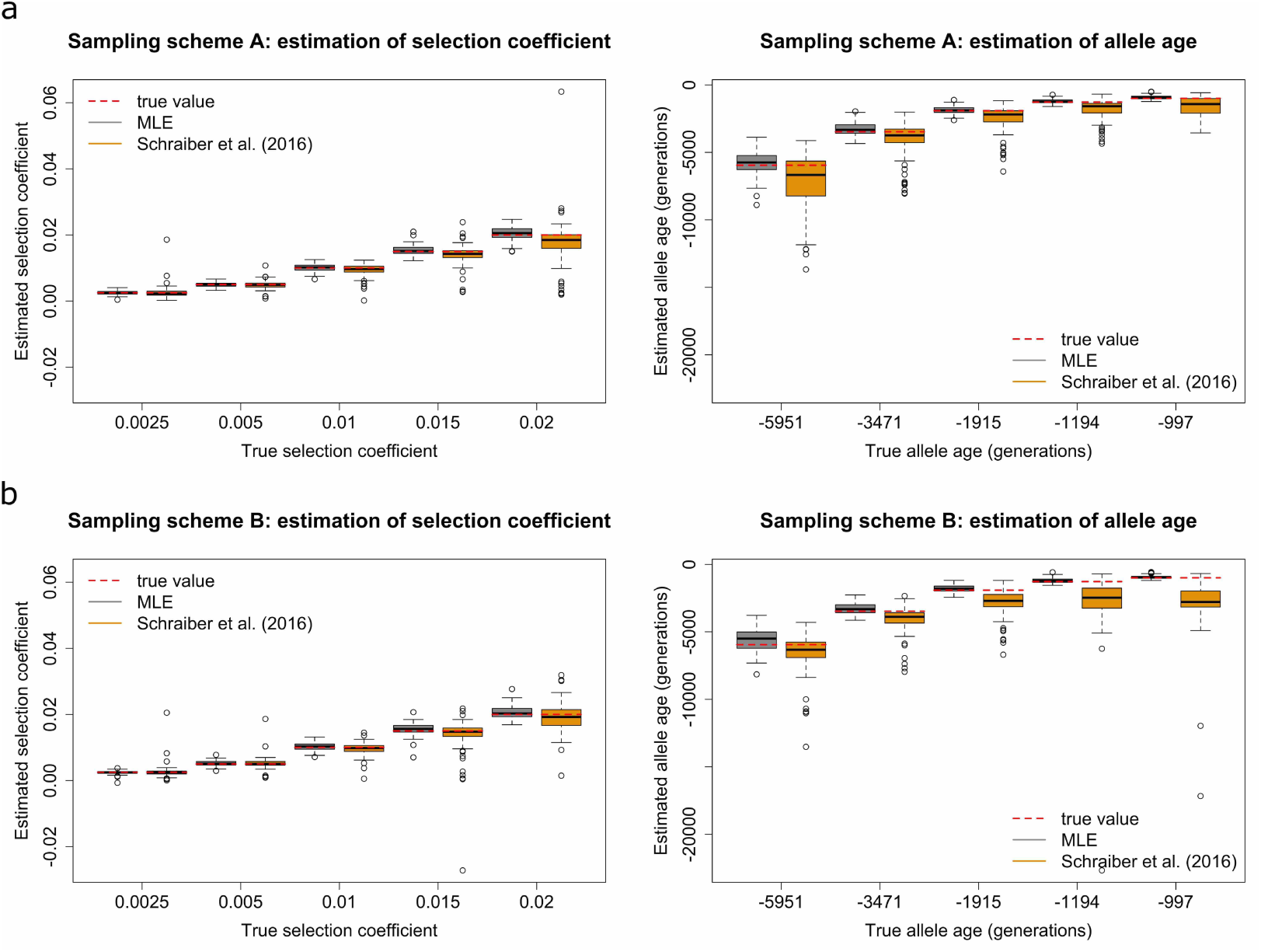
Empirical distributions of estimates for 100 datasets simulated under a constant demographic history with population size *N* (*k*) = 16000 for *k* ∈ {*k*_0_, *k*_0_ + 1, *…*, 0} and dominance parameter *h* = 0.5. Grey boxplots represent the estimates produced with our method, and yellow boxplots using the approach of Schraiber et al. (2016). Boxplots of the estimates for (a) sampling scheme A (b) sampling scheme B.

As shown in Figure 7, the method of Schraiber et al. (2016) performs well for the estimation of the selection coefficient under the constant demographic history when the selection coefficient is small. However, their method shows an increasing trend of underestimation when the selection coefficient is large. Similar results can be found in the performance studies presented in Schraiber et al. (2016). For the allele age, the approach of Schraiber et al. (2016) overestimates its absolute value under the constant demographic history.

In comparison, for all positive selection coefficients, our method performs significantly better in estimating both selection coefficient and allele age under the constant demographic history. Our approach produces reasonably accurate estimates with smaller bias and RMSE across different parameter values and sampling schemes. Non-constant demographic histories cause some deterioration in the performance of our procedure, *i.e.*, larger bias and RMSE in comparison to those under the constant demographic history, but not as much as with the method of Schraiber et al. (2016) (see Supplemental Material, Figures S6-S8 and Tables S12-S15). Indeed, the worst performance of our approach occurs in the case of selective neutrality (*s* = 0) (see Supplemental Material, Figure S9). This can be explained by the following: all else being equal, mutant allele frequency trajectories of the underlying population with larger selection coefficients have less noise and hence more information than smaller selection coefficients.

In summary, for positive selection coefficients, our approach is significantly more accurate across different parameter values, demographic histories and sampling schemes, when compared to the approach of Schraiber et al. (2016). The performance of the method of Schraiber et al. (2016) is also heavily influenced by the demographic history and the sampling scheme.

### 3.2. Analysis of real data

We show the utility of our approach on real data by re-analysing the aDNA data associated with horse coat colouration. Ludwig et al. (2009) sequenced 89 ancient horse samples at eight genes encoding coat colouration, which were obtained from Siberia, Middle and Eastern Europe, China and the Iberian Peninsula, ranging from a pre- to a post-domestication period. Ludwig et al. (2009) applied the method of Bollback et al. (2008) and found two of these genes, *ASIP* and *MC1R*, which showed strong fluctuations in the mutant allele frequencies of the sample, to be likely selectively advantageous. Malaspinas et al. (2012), Steinrücken et al. (2014) and Schraiber et al. (2016) then re-analysed the same aDNA data for *ASIP* and *MC1R* with their approaches incorporating more complex evolutionary scenarios.

We re-analyse the aDNA data for *ASIP* and *MC1R* from Wutke et al. (2016), which contains 107 ancient horse samples sequenced at eight genes encoding coat colouration by them along with 94 sequenced ancient horse samples from the previous studies of Ludwig et al. (2009) and Pruvost et al. (2011). From Der Sarkissian et al. (2015), we take the average length of a generation of the horse to be eight years and show the changes in the mutant allele frequencies of the sample for *ASIP* and *MC1R* through successive generations in Figures 8a and b. As in Schraiber et al. (2016), we apply our method to infer natural selection and allele age for *ASIP* and *MC1R* by explicitly incorporating the horse demographic history of Der Sarkissian et al. (2015), as illustrated in Figure 8c, with the most recent population size *N*_0_ = 16000 being the reference population size. In addition, we set the dominance parameter to *h* = 1 in our analysis as the mutant alleles at the *ASIP* and *MC1R* loci are both recessive (Rieder et al., 2001).

**Figure 8:**
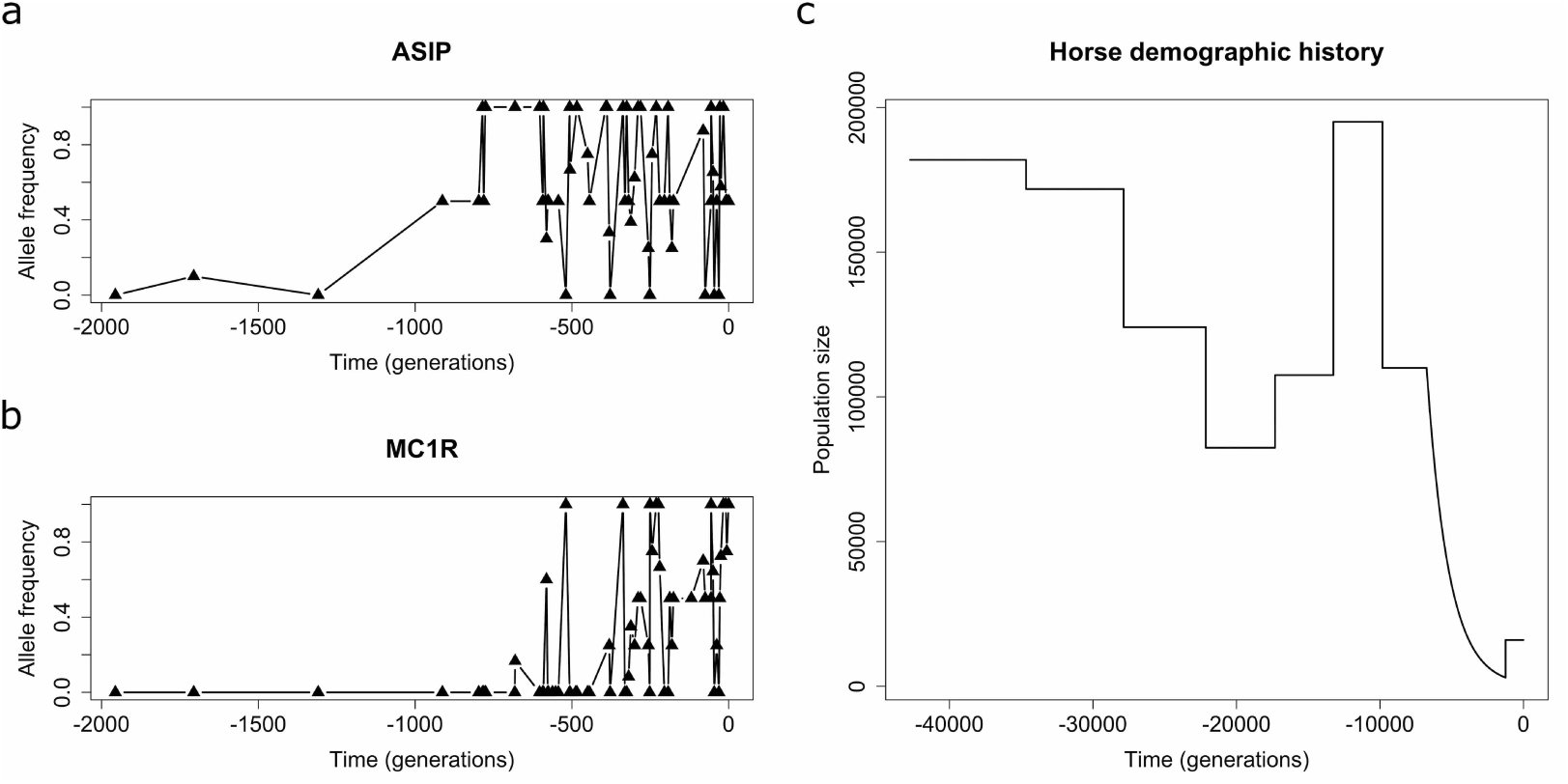
Ancient horse samples sequenced at *ASIP* and *MC1R* from Wutke et al. (2016). Temporal changes in the mutant allele frequencies of the sample for (a) *ASIP* and (b) *MC1R*. (c) Horse demographic history of Der Sarkissian et al. (2015).

We illustrate the resulting likelihood surface for the *ASIP* gene in Figure 9a and find that the likelihood surface attains its maximum at 0.0013 for the selection coefficient and −42982 for the allele age, *i.e.*, 42982 years before present (BP). Our results suggest that the mutation in *ASIP* was favoured by natural selection and was created in the Pleistocene, which lasted from about 2580000 to 11700 years BP (Cohen et al., 2013). To establish the significance of our findings, we compute the corresponding 95% confidence intervals (CI’s) with a bootstrap procedure called case resampling (Efron & Tibshirani, 1994), where we resample with replacement from the 62 time stamped samples in the *ASIP* dataset to form the bootstrap samples. More specifically, we sample with replacement from the 62 sampling time points, which generates exactly 62 time points that may have duplicates. If a certain time point with the observation (*n*_*k*_, *c*_*k*_) is duplicated *m* times in the resampled time points, the bootstrap observation associated with this time point is (*mn*_*k*_, *mc*_*k*_). The 95% CI is [0.0004, 0.0022] for the selection coefficient and [−175272, −18749] for the allele age, respectively, where the 95% CI’s are constructed as the interval between 2.5th and 97.5th percentiles of the estimates for the 1000 bootstrap replicates. This presents further evidence that the mutation in *ASIP* was positively selected and arose in the Middle Pleistocene (lasted from about 773000 to 126000 years BP) or Upper Pleistocene (lasted from about 126000 to 11700 years BP).

**Figure 9:**
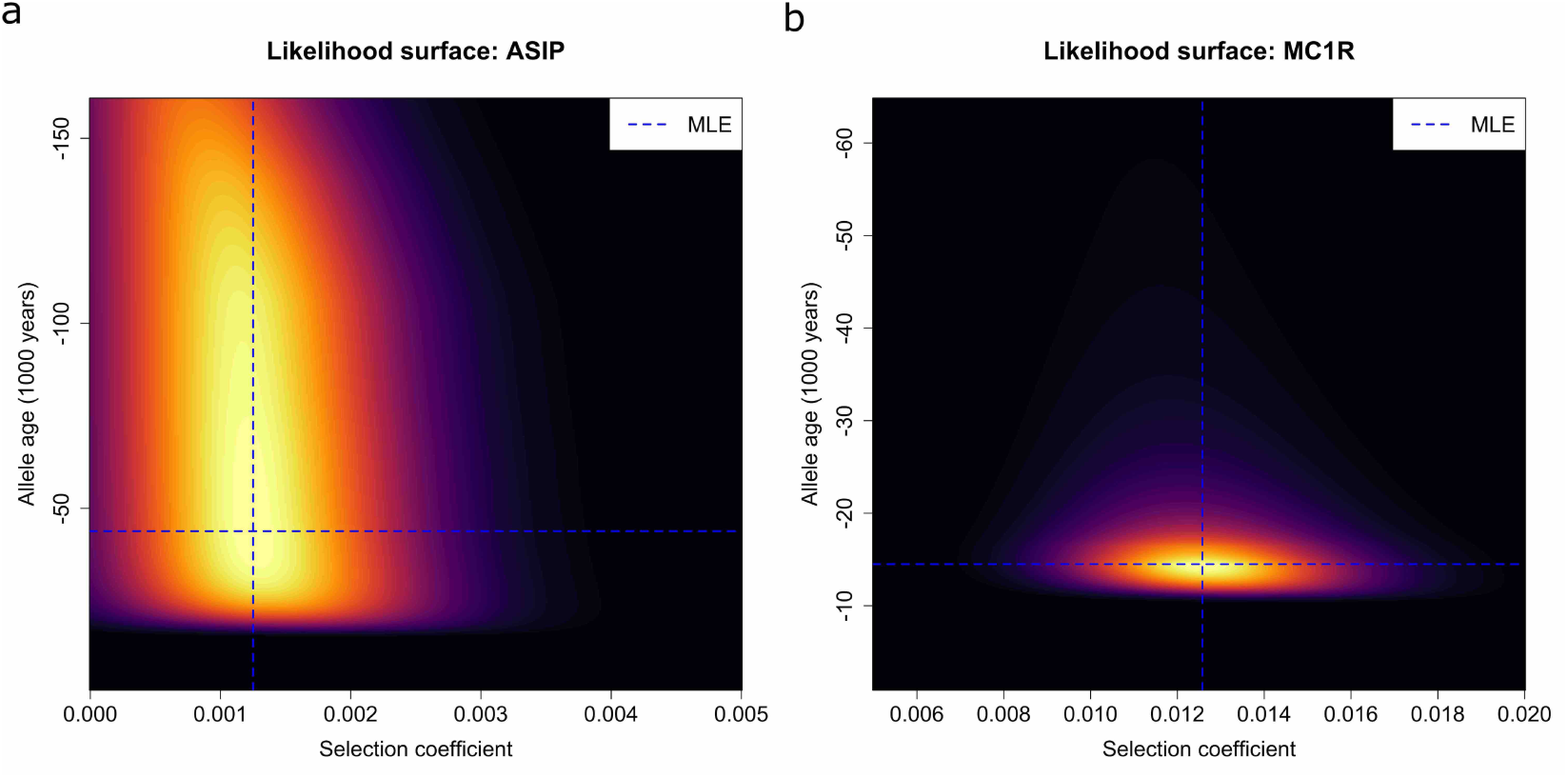
Likelihood surfaces produced under the non-constant demographic history of Der Sarkissian et al. (2015) for (a) *ASIP* and (b) *MC1R*.

The resulting likelihood surface for the *MC1R* gene is illustrated in Figure 9b. Our estimate of the selection coefficient is 0.0126 with the 95% CI [0.0091, 0.0176], which indicates that the mutation in *MC1R* was selectively advantageous. Our estimate of the allele age is −13645 with the 95% CI [−15430, −12866], which suggests that the mutation in *MC1R* was created in the Upper Pleistocene.

It should be noted that one uncertainty our bootstrap procedure does not take into account is that of the underlying demographic history. The demographic history of Der Sarkissian et al. (2015) is an estimate, but we take this to be fixed in our bootstrap procedure. Therefore, the confidence intervals we state above are likely to be narrower than if we have taken into account the further uncertainty in the underlying demographic history.

In summary, our findings suggest that *ASIP* and *MC1R* mutations arose in the Pleistocene and have both been favoured by natural selection. The climate of the Pleistocene saw dramatic changes and included the Last Glacial Maximum (LGM), the coldest phase of the Last Glacial Period (LGP), which lasted from approximately 26500 years BP to 19000 to 20000 years BP (Clark et al., 2009). Our results are weak evidence supporting *ASIP* mutations being created before the LGM (but with probability 0.8), and strong evidence supporting *MC1R* mutations being created after the LGM.

We also present the results under two different constant demographic histories: the first with constant population size 16000, the most recent population size, and the second with constant population size 2500, the bottleneck population size. These results are illustrated in Figures 10 and 11, respectively, with their maximum likelihood estimates and 95% CIs summarised in Supplemental Material, Tables S16. With constant population sizes of both 16000 and 2500, we find strong evidence to support that *ASIP* and *MC1R* mutations are both positively selected. The estimates of the selection coefficient are similar to those produced under the demographic history of Der Sarkissian et al. (2015), but the estimates of the allele age display large variability under different demographic histories. We find that *ASIP* mutations were created during the LGM with constant population size 2500, and *MC1R* mutations were created during the LGM with constant population size 16000, which are in conflict with earlier results obtained under the demographic history of Der Sarkissian et al. (2015). This again demonstrates that changing the underlying demographic history can significantly alter estimates of the allele age.

**Figure 10:**
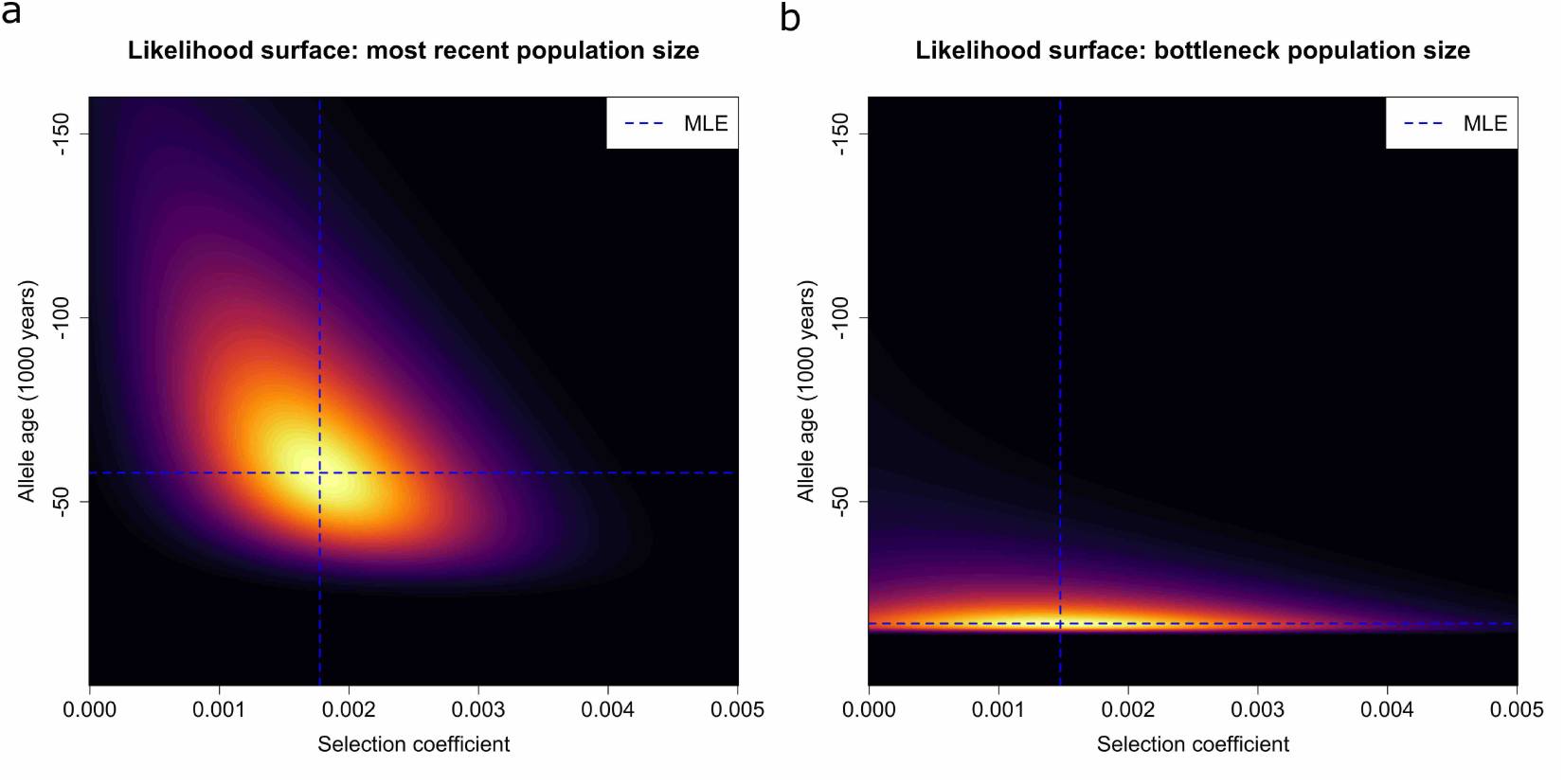
Likelihood surfaces for *ASIP* produced under the constant demographic history with (a) the most recent population size of 16000 and (b) the bottleneck population size of 2500.

**Figure 11:**
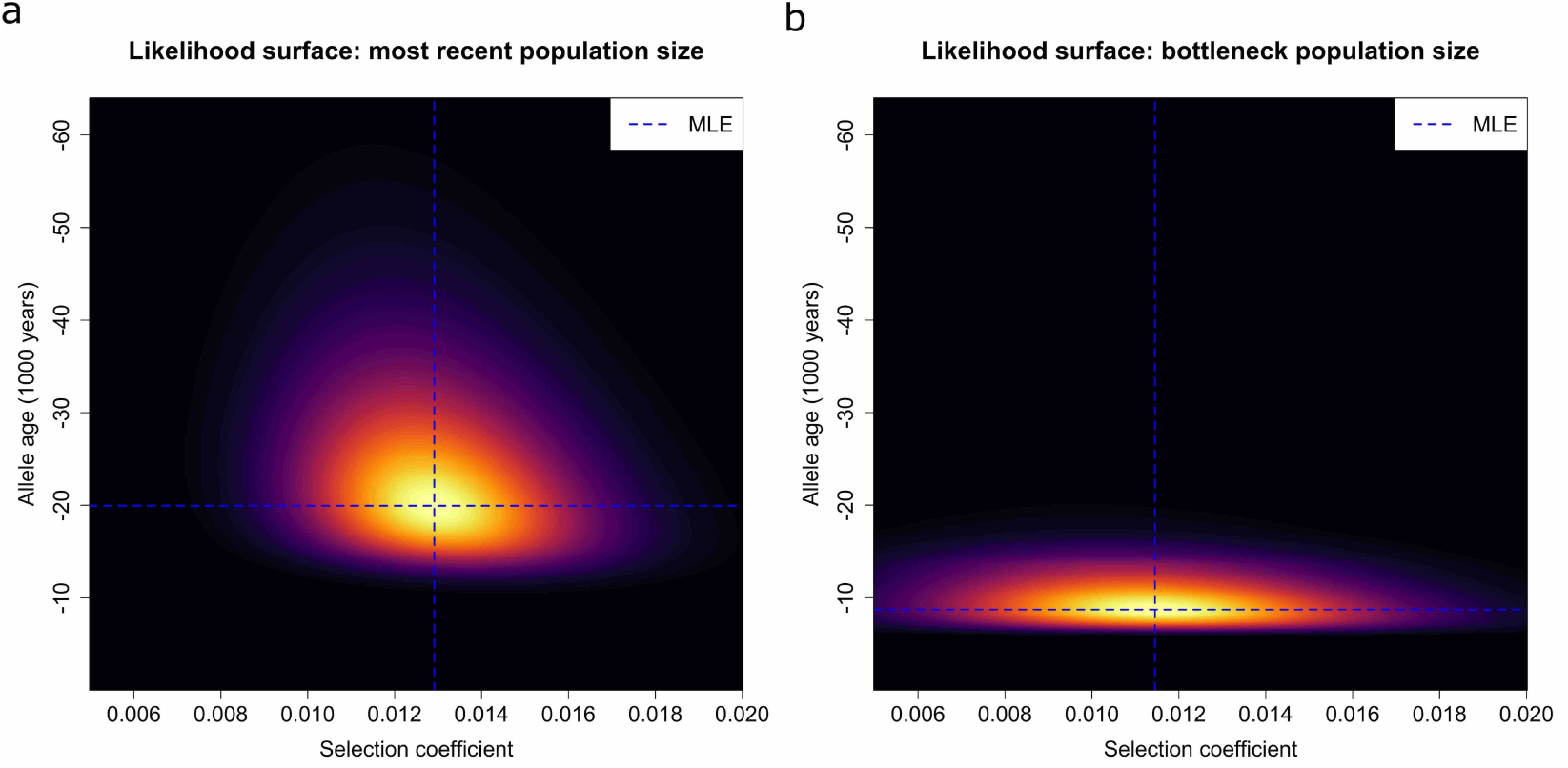
Likelihood surfaces for *MC1R* produced under the constant demographic history with (a) the most recent population size of 16000 and (b) the bottleneck population size of 2500.

### 3.3. Computational issues

We apply the Crank-Nicolson scheme proposed by Crank & Nicolson (1947) to numerically solve the KBE’s in Eqs. (1) and (9). To achieve higher computational accuracy of the numerical solution, it is desirable to increase the number of grid points in the Crank-Nicolson approach, which however becomes burdensome computationally. We adopt a grid that is logarithmically divided close to 0 and 1, and such a partition increases the number of grid points at the two ends of the state space [0, 1]. More specifically, in our analysis we use the grid for the state space [0, 1] that is uniformly divided in [0.03, 0.97] with 30 subintervals and logarithmically divided in [0, 0.03] and [0.97, 1] with 120 subintervals each. To achieve the maximum of the likelihood surface for the selection coefficient and the allele age, we start the section search on the selection coefficient with 7 equally spaced selection coefficients within a fixed interval [−0.02, 0.04].

In our analysis of *ASIP* and *MC1R*, a single run for a fixed value of the selection coefficient takes about 54 seconds for *MC1R* on a single core of an Intel Core i7 processor at 3.5 GHz, whereas the same run takes approximately 84 seconds for *ASIP*. The entire maximum likelihood estimation procedure for *MC1R* takes 905 seconds, whereas this procedure for *ASIP* takes 1728 seconds. All of our code in this work is written in R.

## 4. Discussion

In this work, we developed a novel likelihood-based method for co-estimating selection coefficient and allele age from time series data of allele frequencies under different demographic histories in this work. To our knowledge, Malaspinas et al. (2012) and Schraiber et al. (2016) are the only existing methods that can jointly infer natural selection and allele age from time series data of allele frequencies. Our key innovation is that we take the Wright-Fisher diffusion conditioned to survive until at least the time of the most recent sample to be the underlying process in the HMM framework and calculate the likelihood by numerically solving the KBE resulting from the conditioned Wright-Fisher diffusion backwards in time. For every fixed value of the selection coefficient, we need to numerically solve the KBE only once to obtain the likelihood for all possible values of the allele age. Our backwards-in-time likelihood recursion reduces the two-dimensional numerical search for the maximum of the likelihood surface for the selection coefficient and the allele age, *e.g.*, as used in Malaspinas et al. (2012), to a one-dimensional numerical search over the selection coefficient. Thus, our method can achieve the joint estimates of the selection coefficient and the allele age relatively quickly, *e.g.*, about 1728 and 905 seconds for *ASIP* and *MC1R* from ancient horse samples, respectively. Furthermore, the conditioned Wright-Fisher diffusion incorporated into our HMM framework allows us to avoid the somewhat arbitrary initial condition that the allele is created by mutation at a certain low frequency, *e.g.*, as used in Malaspinas et al. (2012) and Schraiber et al. (2016).

We demonstrated through simulated data that our estimates for both the selection coefficient and the allele age were accurate under a given demographic history while the likelihood surfaces were smooth with a shape that coincided with our intuitive understanding of our approach. Even though the samples are sparsely distributed in time with small uneven sizes, our method still performed well, which is an important feature for aDNA. Our simulation studies also illustrated that ignoring demographic histories had minimal impact on the inference of natural selection but significantly biased the estimation of allele age. In addition, we investigated the impact of the sampling scheme on the inference of natural selection and allele age, showing that within any given sampling period, drawing more chromosomes at fewer time points or the opposite had little effect on the estimation of selection coefficient and allele age. Compared to the method of Schraiber et al. (2016), our approach is superior for the estimation of selection coefficient and allele age in accuracy across different parameter values, demographic histories and sampling schemes.

We re-analysed the genes that have been well studied in previous studies of Ludwig et al. (2009), Malaspinas et al. (2012), Steinrücken et al. (2014) and Schraiber et al. (2016), *ASIP* and *MC1R*, with the expanded aDNA dataset from Wutke et al. (2016). We found strong evidence for weak positive selection acting on the *ASIP* gene and strong positive selection acting on the *MC1R* gene, which was similar to those reported by Ludwig et al. (2009), Steinrücken et al. (2014) and Schraiber et al. (2016). In contrast, Malaspinas et al. (2012) did not have sufficient resolution to distinguish positive selection from negative selection for *ASIP* with their approach from the same aDNA data as analysed in Ludwig et al. (2009). Also, the findings of Malaspinas et al. (2012) suggested that the allele age of *ASIP* mutations ranged from 20000 to 13100 years BP, which was significantly different from our estimate. Such a discrepancy could be caused by different demographic histories, insufficient amounts of data, or improperly grouped samples. Steinrücken et al. (2014) and Schraiber et al. (2016) also inferred the model of natural selection, which we take to be fixed in our analysis with mutant allele homozygotes at both *ASIP* and *MC1R* being recessive. However, thanks to the computational advantages of our approach, it can be readily applied to infer parameters in a model of general diploid natural selection: for example, by running a two-dimensional numerical search over the selection coefficient and the dominance parameter to find the maximum of the likelihood. Our method can also be extended for fluctuating selection coefficients by combining the KBE’s for different population genetic parameters at the change time points suitably. Even though we have only illustrated the utility of our method on aDNA data in this work, our approach can also be used to analyse time series data of allele frequencies from laboratory experiments (*e.g.*, Lang et al., 2013; Wiser et al., 2013; Burke et al., 2014; Le Bihan-Duval et al., 2018; Papkou et al., 2019).

It is worthwhile to note that, unlike earlier studies, where the ancient samples were grouped into 6 sampling time points (*e.g.*, Ludwig et al. (2009)) or 9 sampling time points (*e.g.*, Wutke et al. (2016)), our analysis is conducted on the raw aDNA data, *i.e.*, 201 ancient horse samples from 62 sampling time points. To investigate the effect of grouping samples on the estimates of the selection coefficient and the allele age, we perform an empirical study by random grouping the 62 sampling time points into 9 time points like Wutke et al. (2016) and then apply our method on 1000 randomly grouped samples. As illustrated in Figure 12, the resulting estimates show some variability around the estimates obtained from the raw samples with 62 sampling time points. The relative deviations between the selection coefficient estimated from the grouped samples and the raw samples are 14.35% for *ASIP* and 28.75% for *MC1R*, respectively, and the relative deviations between the allele age estimated from the grouped samples and the raw samples are 35.45% for *ASIP* and 7.57% for *MC1R*, respectively, which implies that grouping samples can significantly alter the estimates of the selection coefficient and the allele age.

**Figure 12:**
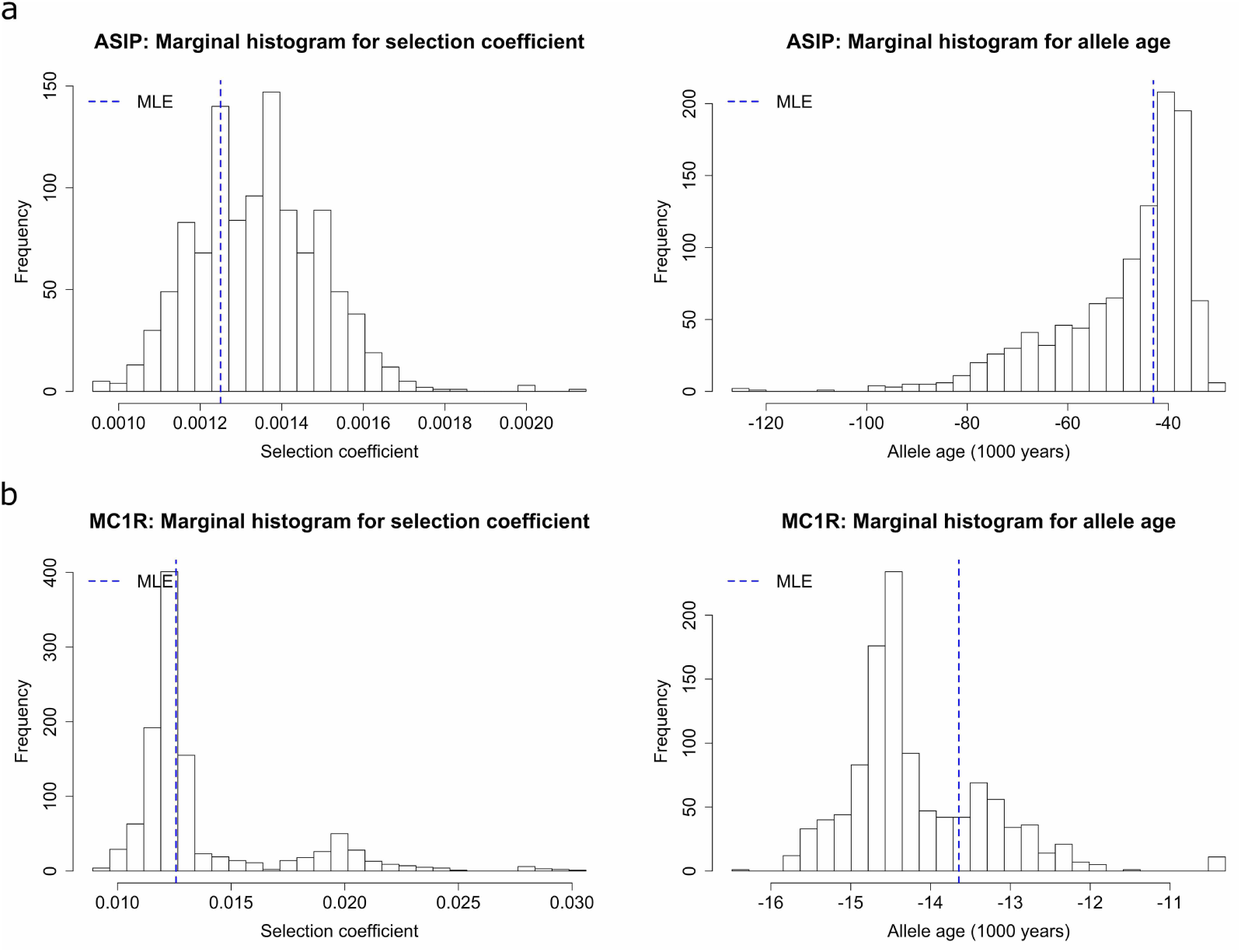
Effects of grouping samples on the estimates of the selection coefficient and the allele age. Marginal histograms of the estimates for 1000 randomly grouped datasets of the temporally spaced samples for (a) *ASIP* and (b) *MC1R*.

Recent findings of Sandoval-Castellanos et al. (2017) suggested that at the end of the Ice Age forests started to advance in Europe, and horses became increasingly dark in coat colour as they left the plains and adapted to live in forests, where dark-coloured coats would have helped them to hide. Our approach does not provide evidence of the onset of natural selection. It delivers estimates of the selection coefficient and the allele age. However, one possibility that is consistent with our results is that horse base coat colour variation could be associated with their adaptation to the transition from a glacial period to an interglacial period. Polymorphisms at the *ASIP* and *MC1R* loci are associated with horse base coat colouration, creating a bay, black or chestnut coat (Rieder et al., 2001), *i.e., ASIP* mutations give rise to black-coloured horses, whereas *MC1R* mutations give rise to chestnut-coloured horses. From Section 3.2, our 95% confidence interval for the allele age of *ASIP* mutations include the LGM but lies mostly before the LGM. This raises the possibility that *ASIP* mutations, giving dark horses, might have been created before the LGM, whereas *MC1R* mutations, leading to paler horses, were created after the LGM. Given the findings of Finch et al. (1984) that dark-coated animals absorb heat more rapidly from solar radiation than light-coated ones, we speculate that *ASIP* mutations were favoured in horses living in cold environments during the glacial period, where dark coats would have helped them to survive, especially for the LGM. In contrast, *MC1R* mutations were favoured when the climate warmed from the late Pleistocene to the early Holocene since it might be then advantageous to be light-coated in warm environments for horses.

One key limitation of our method is that we infer natural selection and allele age without accounting for the interactions between the genes. Such interactions can be epistatic interaction, genetic linkage or others, *e.g.*, the *ASIP* and *MC1R* genes are found with epistatic interactions (Rieder et al., 2001), which may affect population genetic analyses. With the increasing availability of aDNA data across multiple loci, performing accurate estimation on relevant population genetic quantities of interest while accounting for interactions among loci becomes more and more important. To extend our method to allele frequency time series data across multiple loci subject to epistatic interaction or genetic linkage, a significant challenge would be to find an alternative to calculate transition probabilities by numerically solving the KBE in our likelihood computation since it is computationally challenging and prohibitively expensive to numerically solve a high-dimensional partial differential equation.

## Acknowledgements

We would like to thank the anonymous reviewers and the communicating editors for improving this work with their helpful comments. This work was funded in part by the Engineering and Physical Sciences Research Council (EPSRC) Grant EP/I028498/1 to F.Y.

## Notes

### Competing Interest Statement

The authors have declared no competing interest.

### Summary of Updates

We added a paragraph that explains the uncertainty of the estimated demographic history and that our confidence intervals are likely to be narrower than if this uncertainly is taken into account in our real data analyses. We shortened the manuscript and corrected the typos.

