## Supplemental Material for "Maximum likelihood estimation of natural selection and allele age from time series data of allele frequencies"

### File S1. Boundary condition for the conditioned Wright-Fisher diffusion

One problem that arises when numerically solving the Kolmogorov backward equation (KBE) associated with the the conditioned Wright-Fisher diffusion $X^*$ is that $P_\tau(t, 0) = 0$, which means that $\log P_\tau(t, 0)$ needed for the evaluation of $a^*(t, x)$ is $-\infty$. Therefore, we can expect $\frac{\partial}{\partial x} \log P_\tau(t, x)$ to blow up to $\infty$ as $x$ approaches 0. However, $b(t, x)^2 = x(1-x)/\rho(t)^2$, which is roughly $x/\rho(t)^2$ when $x$ is small. Indeed, we shall see that $\lim_{x \downarrow 0} x \frac{\partial}{\partial x} \log P_\tau(t, x) = 1$ for $t \ll \tau$. By Eq. (6.18) of Karlin & Taylor (1981), the conditioned Wright-Fisher diffusion $X^*$ has an entrance boundary at 0, *i.e.*, the conditioned Wright-Fisher diffusion $X^*$ starts at 0 but reaches positive values immediately after that and will never return to 0.

Under the Wright-Fisher diffusion

$$dX(t) = \alpha X(t)(1 - X(t))((1-h) - (1-2h)X(t))dt + \sqrt{\frac{X(t)(1-X(t))}{\rho(t)}} dW(t), \quad t \geq t_0, \quad (1)$$

the fixation probability, *i.e.*, the probability of the Wright-Fisher diffusion $X$ reaching 1 eventually, is $\mathcal{O}(1)$ for large $N$ if $X(t_0) = 1/(2N)$, *i.e.*, this probability does not go to 0 as $N \to \infty$. If $X$ reaches levels significantly larger than $1/(2N)$, in other words, if the mutant allele becomes *established* in the population, then the probability of eventual fixation becomes near certain. Therefore, the key to determining whether the diffusion process reaches fixation eventually is in the early stage of the selective sweep, *i.e.*, when $X$ is small. We assume that $\rho(t)$ is roughly constant over the time that it takes the mutant allele to become established, which is $\mathcal{O}(\log N/N)$. This simplifies the calculation that follows. In this parameter regime, we can approximate the evolution of the Wright-Fisher diffusion in Eq. (1) with a diffusion process $Y_1$ in the case of $h \neq 1$:

$$dY_1(t) = \alpha(1-h)Y_1(t)dt + \sqrt{\frac{Y_1(t)}{\rho}} dW(t), \quad t \geq t_0, \quad (2)$$

and a diffusion process $Y_2$ in the case of $h = 1$:

$$dY_2(t) = \alpha Y_2(t)^2 dt + \sqrt{\frac{Y_2(t)}{\rho}} dW(t), \quad t \geq t_0. \quad (3)$$

The probability that $X$ becomes established is very close to $Y_1$ or $Y_2$ becoming established. Furthermore, once $Y_1$ or $Y_2$ becomes established, since $\alpha = \mathcal{O}(2N)$, *i.e.*, the effect of the drift term is much larger than that of the diffusion term in either Eq. (2) or (3), it is almost certain to go to $\infty$.

The probability of a diffusion process reaching a level $c$ before hitting 0 can be calculated using the speed and scale functions (see *e.g.*, Karlin & Taylor (1981)). Let $T_x$ denote the first hitting time of $x$ and $\mathbb{P}^x$ denote the probability measure of the diffusion process started at $x$. For the diffusion process $Y_1$ in Eq. (2), we have

$$S(x) = \int_0^x s(\xi)\, d\xi = \frac{1 - e^{-\alpha(1-h)\rho x}}{\alpha(1-h)\rho},$$

where

$$s(\xi) = e^{-\alpha(1-h)\rho\xi}.$$

Then the probability of $Y_1$ hitting $c$ before hitting 0 is

$$\mathbb{P}^x\left(T_c < T_0\right) = \frac{S(x) - S(0)}{S(c) - S(0)} = \frac{1 - e^{-\alpha(1-h)\rho x}}{1 - e^{-\alpha(1-h)\rho c}}.$$

Taking $c \to \infty$ yields

$$\mathbb{P}^x\left(T_\infty < T_0\right) = 1 - e^{-\alpha(1-h)\rho x}.$$

As discussed earlier, for $t \ll \tau$ and $x$ close 0, $\mathbb{P}^x\left(T_\infty < T_0\right)$ is very close to the value of $P_\tau(t, x)$. We then have

$$\begin{aligned}\lim_{x\downarrow 0} x\frac{\partial}{\partial x}\log P_\tau(t,x) &\approx \lim_{x\downarrow 0} x\frac{\partial}{\partial x}\log\left(1 - e^{-\alpha(1-h)\rho x}\right)\\ &= \lim_{x\downarrow 0} x\frac{\alpha(1-h)\rho e^{-\alpha(1-h)\rho x}}{1 - e^{-\alpha(1-h)\rho x}}\\ &= 1. \qquad (4)\end{aligned}$$

Similarly, for the diffusion process $Y_2$ in Eq. (3), we have

$$S(x) = \int_0^x s(\xi)\, d\xi = \sqrt{\frac{2\pi}{\alpha\rho}}\mathbb{P}\left(|\mathcal{N}(0,1)| < \sqrt{\alpha\rho}x\right),$$

where

$$s(\xi) = e^{-\frac{\alpha}{2}\rho\xi^2}$$

and $\mathcal{N}(0,1)$ denotes a standard normal random variable. Then the probability of $Y_2$ hitting $c$ before hitting 0 is

$$\mathbb{P}^x(T_c < T_0) = \frac{S(x)-S(0)}{S(c)-S(0)} = \frac{\mathbb{P}\left(|\mathcal{N}(0,1)| < \sqrt{\alpha\rho}x\right)}{\mathbb{P}\left(|\mathcal{N}(0,1)| < \sqrt{\alpha\rho}c\right)},$$

Taking $c \to \infty$ yields

$$\mathbb{P}^x\left(T_\infty < T_0\right) = \mathbb{P}\left(|\mathcal{N}(0,1)| < \sqrt{\alpha\rho}x\right).$$

This implies

$$\begin{aligned}\lim_{x\downarrow 0} x\frac{\partial}{\partial x}\log P_\tau(t,x) &\approx \lim_{x\downarrow 0} x\frac{\partial}{\partial x}\log \mathbb{P}\left(|\mathcal{N}(0,1)| < \sqrt{\alpha\rho}x\right)\\ &= \lim_{x\downarrow 0} x\frac{\partial}{\partial x}\log\left(\sqrt{\frac{2\alpha\rho}{\pi}}x + \mathcal{O}(x^2)\right)\\ &= \lim_{x\downarrow 0} x\frac{\sqrt{\frac{2\alpha\rho}{\pi}} + \mathcal{O}(x)}{\sqrt{\frac{2\alpha\rho}{\pi}}x + \mathcal{O}(x^2)}\\ &= 1. \end{aligned} \tag{5}$$

Combining Eqs. (4) and (5), we have

$$\lim_{x\downarrow 0} x\frac{\partial}{\partial x}\log P_\tau(t,x) = 1$$

for $t \ll \tau$.

### File S2. Additional results on the analysis of simulated data

| Selection coefficient | | | Allele age | | |
|---|---|---|---|---|---|
| True value | Bias | RMSE | True value | Bias | RMSE |
| 0.000e-00 | 3.480e-04 | 1.053e-03 | -6.999e+03 | 1.127e+03 | 4.819e+03 |
| 2.500e-03 | -1.710e-04 | 5.790e-04 | -6.267e+03 | -1.997e+02 | 5.033e+02 |
| 5.000e-03 | -2.360e-04 | 9.020e-04 | -3.439e+03 | -1.550e+02 | 3.425e+02 |
| 1.000e-02 | -2.910e-04 | 1.598e-03 | -1.778e+03 | -3.599e+01 | 1.657e+02 |
| 1.500e-02 | -4.970e-04 | 1.865e-03 | -1.262e+03 | -2.841e+01 | 1.048e+02 |
| 2.000e-02 | -5.510e-04 | 2.479e-03 | -9.410e+02 | -2.859e+01 | 7.980e+01 |

(a) Sampling scheme A: 20 chromosomes drawn at 60 sampling time points.

| Selection coefficient | | | Allele age | | |
|---|---|---|---|---|---|
| True value | Bias | RMSE | True value | Bias | RMSE |
| 0.000e-00 | 3.290e-04 | 9.170e-04 | -6.999e+03 | 1.689e+03 | 6.409e+03 |
| 2.500e-03 | -1.240e-04 | 5.570e-04 | -6.267e+03 | -2.446e+02 | 5.315e+02 |
| 5.000e-03 | -1.810e-04 | 8.320e-04 | -3.439e+03 | -1.170e+02 | 3.094e+02 |
| 1.000e-02 | -4.180e-04 | 1.469e-03 | -1.778e+03 | -4.812e+01 | 1.464e+02 |
| 1.500e-02 | -5.330e-04 | 2.009e-03 | -1.262e+03 | -2.412e+01 | 1.067e+02 |
| 2.000e-02 | -5.290e-04 | 2.271e-03 | -9.410e+02 | -2.085e+01 | 7.180e+01 |

(b) Sampling scheme B: 120 chromosomes drawn at 10 sampling time points.

Table S1: Bias and RMSE of estimates for 100 datasets simulated under a constant demographic history with population size $N(k) = 16000$ for $k \in \{k_0, k_0 + 1, \dots, 0\}$ and dominance parameter $h = 0$.

| Selection coefficient | | | Allele age | | |
|---|---|---|---|---|---|
| True value | Bias | RMSE | True value | Bias | RMSE |
| 0.000e-00 | 3.480e-04 | 1.053e-03 | -6.999e+03 | 1.127e+03 | 4.819e+03 |
| 2.500e-03 | -8.000e-06 | 6.320e-04 | -5.951e+03 | -1.698e+02 | 8.620e+02 |
| 5.000e-03 | 3.100e-05 | 6.670e-04 | -3.471e+03 | -2.240e+02 | 5.597e+02 |
| 1.000e-02 | -3.900e-05 | 1.148e-03 | -1.915e+03 | -5.498e+01 | 2.828e+02 |
| 1.500e-02 | -3.290e-04 | 1.408e-03 | -1.277e+03 | -6.539e+01 | 1.772e+02 |
| 2.000e-02 | -5.500e-04 | 2.101e-03 | -9.970e+02 | -7.842e+01 | 1.679e+02 |

(a) Sampling scheme A: 20 chromosomes drawn at 60 sampling time points.

| Selection coefficient | | | Allele age | | |
|---|---|---|---|---|---|
| True value | Bias | RMSE | True value | Bias | RMSE |
| 0.000e-00 | 3.290e-04 | 9.170e-04 | -6.999e+03 | 1.689e+03 | 6.409e+03 |
| 2.500e-03 | 5.800e-05 | 5.920e-04 | -5.951e+03 | -3.339e+02 | 9.067e+02 |
| 5.000e-03 | -2.100e-04 | 7.980e-04 | -3.471e+03 | -1.717e+02 | 4.354e+02 |
| 1.000e-02 | -2.080e-04 | 1.203e-03 | -1.915e+03 | -1.272e+02 | 2.899e+02 |
| 1.500e-02 | -5.400e-04 | 1.771e-03 | -1.277e+03 | -8.075e+01 | 1.950e+02 |
| 2.000e-02 | -6.360e-04 | 1.941e-03 | -9.970e+02 | -4.776e+01 | 1.461e+02 |

(b) Sampling scheme B: 120 chromosomes drawn at 10 sampling time points.

Table S2: Bias and RMSE of estimates for 100 datasets simulated under a constant demographic history with population size $N(k) = 16000$ for $k \in \{k_0, k_0 + 1, \dots, 0\}$ and dominance parameter $h = 0.5$.

| Selection coefficient | | | Allele age | | |
|---|---|---|---|---|---|
| True value | Bias | RMSE | True value | Bias | RMSE |
| 0.000e-00 | 3.480e-04 | 1.053e-03 | -6.999e+03 | 1.127e+03 | 4.819e+03 |
| 2.500e-03 | 6.760e-04 | 3.084e-03 | -1.283e+04 | -5.217e+02 | 1.887e+03 |
| 5.000e-03 | 1.847e-03 | 5.234e-03 | -7.373e+03 | -1.393e+02 | 2.055e+03 |
| 1.000e-02 | 2.135e-03 | 7.134e-03 | -5.736e+03 | -1.858e+02 | 1.498e+03 |
| 1.500e-02 | 2.111e-03 | 8.568e-03 | -4.526e+03 | -2.558e+02 | 1.008e+03 |
| 2.000e-02 | 4.173e-03 | 1.315e-02 | -3.902e+03 | -9.842e+01 | 9.935e+02 |

(a) Sampling scheme A: 20 chromosomes drawn at 60 sampling time points.

| Selection coefficient | | | Allele age | | |
|---|---|---|---|---|---|
| True value | Bias | RMSE | True value | Bias | RMSE |
| 0.000e-00 | 3.290e-04 | 9.170e-04 | -6.999e+03 | 1.689e+03 | 6.409e+03 |
| 2.500e-03 | -4.000e-05 | 8.830e-04 | -1.283e+04 | -3.554e+02 | 1.881e+03 |
| 5.000e-03 | 2.343e-03 | 6.363e-03 | -7.373e+03 | -3.769e+02 | 1.549e+03 |
| 1.000e-02 | 5.600e-04 | 4.925e-03 | -5.736e+03 | -4.655e+02 | 1.030e+03 |
| 1.500e-02 | 3.330e-03 | 9.964e-03 | -4.526e+03 | -2.131e+02 | 9.326e+02 |
| 2.000e-02 | 5.342e-03 | 1.525e-02 | -3.902e+03 | -1.084e+02 | 1.269e+03 |

(b) Sampling scheme B: 120 chromosomes drawn at 10 sampling time points.

Table S3: Bias and RMSE of estimates for 100 datasets simulated under a constant demographic history with population size $N(k) = 16000$ for $k \in \{k_0, k_0 + 1, \ldots, 0\}$ and dominance parameter $h = 1$.

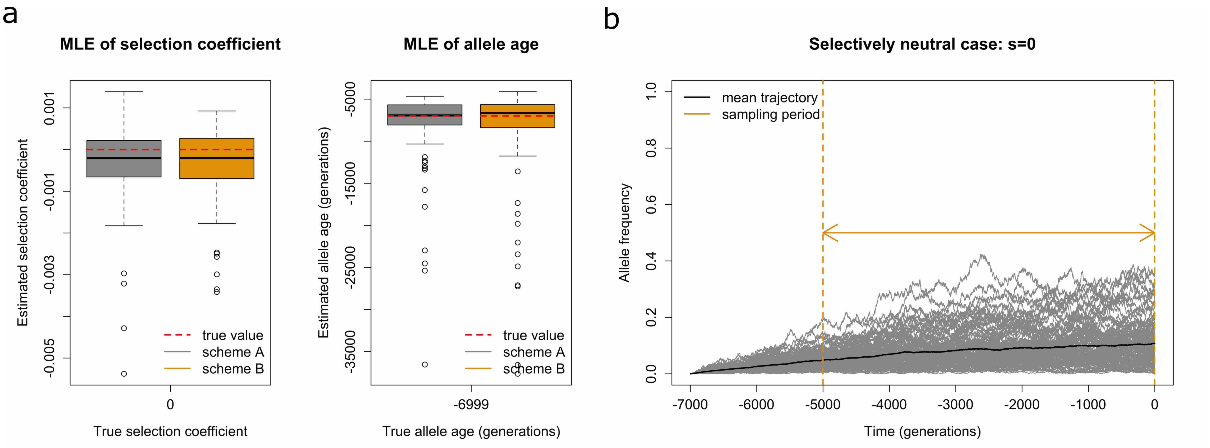


Figure S1: Empirical studies of a *selectively neutral* locus under a constant demographic history with population size $N(k) = 16000$ for $k \in \{k_0, k_0 + 1, \ldots, 0\}$ and selection coefficient $s = 0$. (a) Boxplots of estimates for 100 simulated datasets. (b) Simulated mutant allele frequency trajectories of the underlying population.

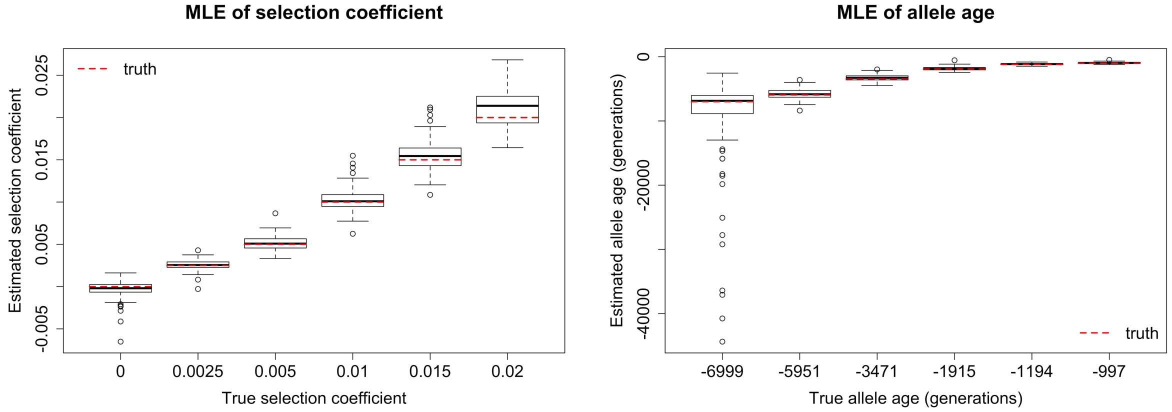


Figure S2: Empirical distributions of estimates for 100 datasets simulated under a constant demographic history with population size $N(k) = 16000$ for $k \in \{k_0, k_0 + 1, \ldots, 0\}$ and dominance parameter $h = 0.5$. The sampling time points are the same as those in sampling scheme B, *i.e.*, 10 sampling points. The sample sizes are independent Poisson distributed with parameter 60.

| Selection coefficient | | | Allele age | | |
|---|---|---|---|---|---|
| True value | Bias | RMSE | True value | Bias | RMSE |
| 0.000e-00 | 3.920e-04 | 1.156e-03 | -6.999e+03 | 2.527e+03 | 8.053e+03 |
| 2.500e-03 | -5.800e-05 | 6.070e-04 | -5.951e+03 | -1.610e+02 | 8.493e+02 |
| 5.000e-03 | -1.200e-04 | 8.420e-04 | -3.471e+03 | -1.967e+02 | 5.246e+02 |
| 1.000e-02 | -2.830e-04 | 1.432e-03 | -1.915e+03 | -7.826e+01 | 2.936e+02 |
| 1.500e-02 | -4.970e-04 | 1.925e-03 | -1.194e+03 | -4.149e+01 | 1.543e+02 |
| 2.000e-02 | -1.043e-03 | 2.508e-03 | -9.970e+02 | -3.918e+01 | 1.323e+02 |

Table S4: Bias and RMSE of estimates for 100 datasets simulated under a constant demographic history with population size $N(k) = 16000$ for $k \in \{k_0, k_0 + 1, \ldots, 0\}$ and dominance parameter $h = 0.5$. The sampling time points are the same as those in sampling scheme B, *i.e.*, 10 sampling points. The sample sizes are independent Poisson distributed with parameter 60.

| Selection coefficient | | | Allele age | | |
|---|---|---|---|---|---|
| True value | Bias | RMSE | True value | Bias | RMSE |
| 0.000e-00 | 3.100e-04 | 9.250e-04 | -6.999e+03 | 1.861e+03 | 6.701e+03 |
| 2.500e-03 | 2.900e-05 | 6.550e-04 | -5.951e+03 | -2.726e+02 | 9.584e+02 |
| 5.000e-03 | -6.900e-05 | 9.580e-04 | -3.471e+03 | -1.175e+02 | 4.598e+02 |
| 1.000e-02 | 6.000e-05 | 1.213e-03 | -1.915e+03 | -8.090e+01 | 2.627e+02 |
| 1.500e-02 | -5.310e-04 | 1.832e-03 | -1.194e+03 | -5.766e+01 | 1.757e+02 |
| 2.000e-02 | -2.960e-04 | 1.909e-03 | -9.970e+02 | -3.544e+01 | 1.381e+02 |

(a) Sampling scheme A: 20 chromosomes drawn at 60 sampling time points.

| Selection coefficient | | | Allele age | | |
|---|---|---|---|---|---|
| True value | Bias | RMSE | True value | Bias | RMSE |
| 0.000e-00 | 3.330e-04 | 9.520e-04 | -6.999e+03 | 1.769e+03 | 6.168e+03 |
| 2.500e-03 | 2.900e-05 | 6.240e-04 | -5.951e+03 | -2.916e+02 | 9.074e+02 |
| 5.000e-03 | -1.040e-04 | 9.170e-04 | -3.471e+03 | -1.282e+02 | 4.435e+02 |
| 1.000e-02 | 1.030e-04 | 1.334e-03 | -1.915e+03 | -6.538e+01 | 2.672e+02 |
| 1.500e-02 | -3.270e-04 | 1.565e-03 | -1.194e+03 | -4.735e+01 | 1.677e+02 |
| 2.000e-02 | -3.970e-04 | 2.100e-03 | -9.970e+02 | -3.240e+01 | 1.306e+02 |

(b) Sampling scheme B: 120 chromosomes drawn at 10 sampling time points.

Table S5: Bias and RMSE of estimates for 100 datasets simulated under a bottleneck demographic history with population size $N(k) = 8000$ for $k \in \{[k_0/2], [k_0/2]+1, \dots, [3k_0/4]\}$ and $N(k) = 16000$ otherwise, and dominance parameter $h = 0.5$. The estimates are obtained assuming true demographic history.

| Selection coefficient | | | Allele age | | |
|---|---|---|---|---|---|
| True value | Bias | RMSE | True value | Bias | RMSE |
| 0.000e-00 | 9.130e-04 | 1.573e-03 | -6.999e+03 | -3.850e+02 | 2.766e+03 |
| 2.500e-03 | 7.700e-05 | 7.430e-04 | -5.951e+03 | -7.693e+02 | 1.101e+03 |
| 5.000e-03 | -1.500e-05 | 9.930e-04 | -3.471e+03 | -3.829e+02 | 5.549e+02 |
| 1.000e-02 | 1.000e-04 | 1.214e-03 | -1.915e+03 | -2.160e+02 | 3.201e+02 |
| 1.500e-02 | -4.710e-04 | 1.832e-03 | -1.194e+03 | -1.444e+02 | 2.111e+02 |
| 2.000e-02 | -2.030e-04 | 1.938e-03 | -9.970e+02 | -1.046e+02 | 1.642e+02 |

(a) Sampling scheme A: 20 chromosomes drawn at 60 sampling time points.

| Selection coefficient | | | Allele age | | |
|---|---|---|---|---|---|
| True value | Bias | RMSE | True value | Bias | RMSE |
| 0.000e-00 | 9.290e-04 | 1.608e-03 | -6.999e+03 | -3.369e+02 | 2.548e+03 |
| 2.500e-03 | 9.300e-05 | 7.510e-04 | -5.951e+03 | -7.663e+02 | 1.064e+03 |
| 5.000e-03 | -3.400e-05 | 9.450e-04 | -3.471e+03 | -3.810e+02 | 5.380e+02 |
| 1.000e-02 | 1.420e-04 | 1.329e-03 | -1.915e+03 | -2.015e+02 | 3.121e+02 |
| 1.500e-02 | -3.010e-04 | 1.577e-03 | -1.194e+03 | -1.380e+02 | 2.040e+02 |
| 2.000e-02 | -3.370e-04 | 2.120e-03 | -9.970e+02 | -1.033e+02 | 1.588e+02 |

(b) Sampling scheme B: 120 chromosomes drawn at 10 sampling time points.

Table S6: Bias and RMSE of estimates for 100 datasets simulated under a bottleneck demographic history with population size $N(k) = 8000$ for $k \in \{[k_0/2], [k_0/2]+1, \dots, [3k_0/4]\}$ and $N(k) = 16000$ otherwise, and dominance parameter $h = 0.5$. The estimates are obtained assuming a constant demographic history with population size $N(k) = 8000$ for $k \in \{k_0, k_0 + 1, \dots, 0\}$.

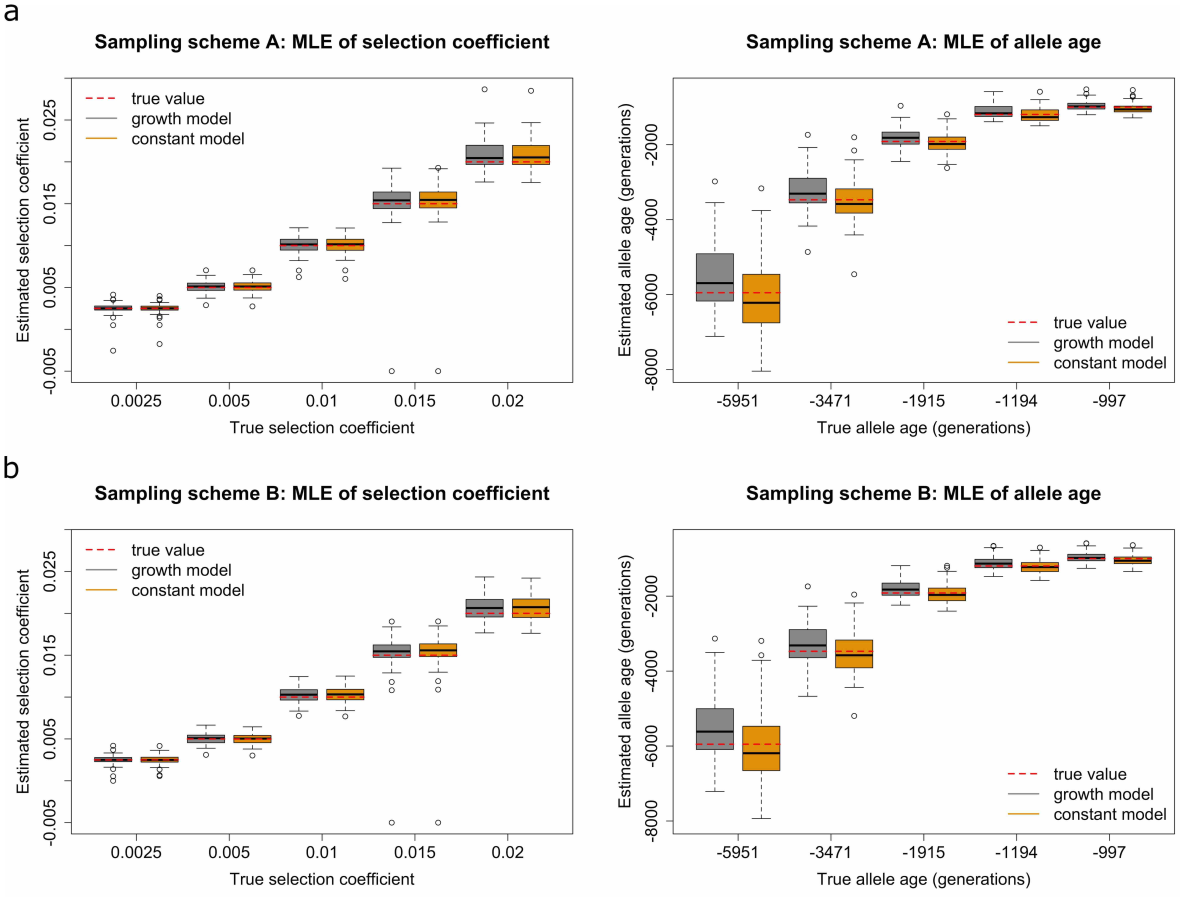


Figure S3: Empirical distributions of estimates for 100 datasets simulated under a growth demographic history with population size $N(k) = 16000$ for $k \in \{k_0, k_0 + 1, \ldots, [k_0/2]\}$ and $N(k) = 32000$ otherwise, and dominance parameter $h = 0.5$. Grey boxplots represent estimates produced assuming the true demographic history, and yellow boxplots assuming a constant demographic history with the most recent population size $N(k) = 32000$ for $k \in \{k_0, k_0 + 1, \ldots, 0\}$. Boxplots of the estimates for (a) sampling scheme A (b) sampling scheme B.

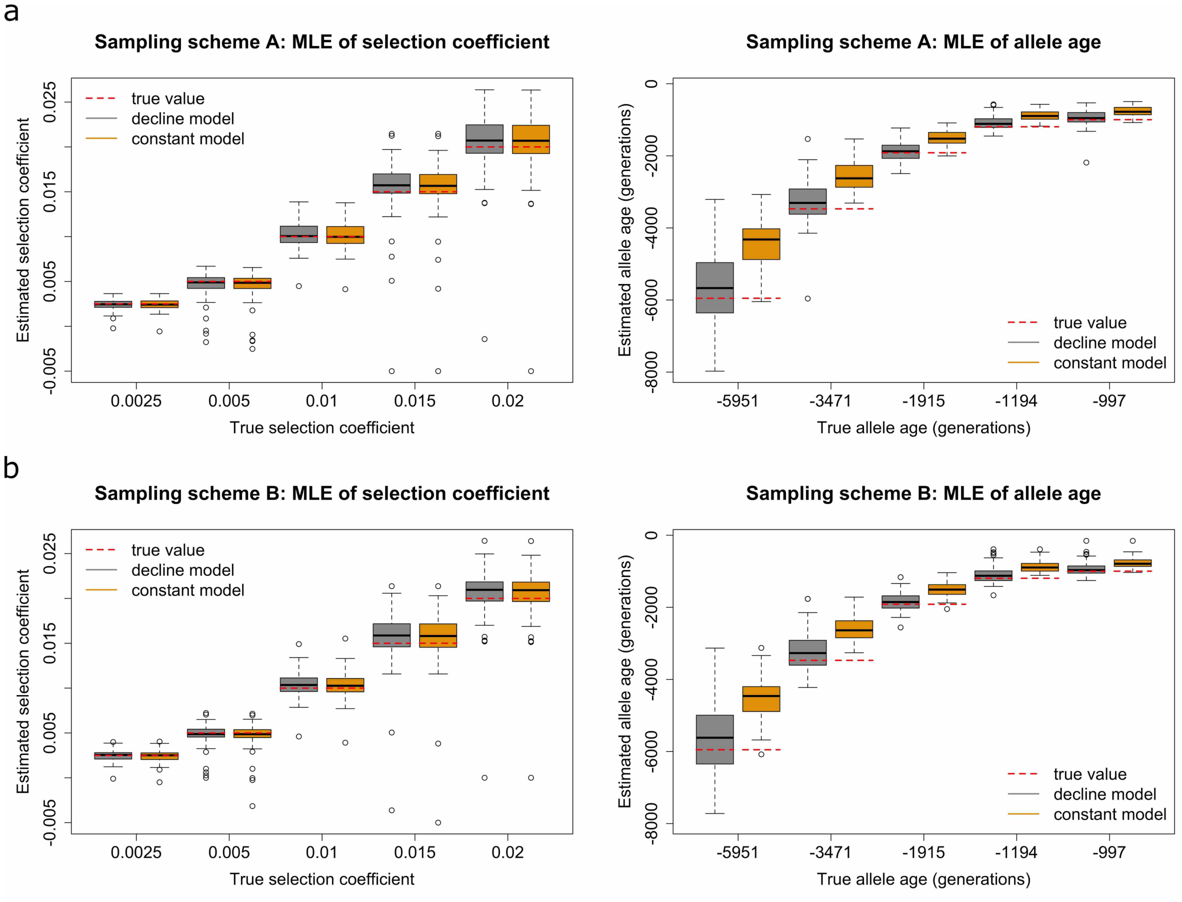


Figure S4: Empirical distributions of estimates for 100 datasets simulated under a decline demographic history with population size $N(k) = 100000$ for $k \in \{k_0, k_0+1, \ldots, [k_0/2]\}$ and $N(k) = 16000$ otherwise, and dominance parameter $h = 0.5$. Grey boxplots represent estimates produced assuming the true demographic history, and yellow boxplots assuming a constant demographic history with the most recent population size $N(k) = 16000$ for $k \in \{k_0, k_0+1, \ldots, 0\}$. Boxplots of the estimates for (a) sampling scheme A (b) sampling scheme B.

| Selection coefficient | | | Allele age | | |
|---|---|---|---|---|---|
| True value | Bias | RMSE | True value | Bias | RMSE |
| 0.000e-00 | 4.290e-04 | 1.058e-03 | -6.999e+03 | 1.821e+03 | 5.655e+03 |
| 2.500e-03 | 1.700e-05 | 6.870e-04 | -5.951e+03 | -4.304e+02 | 9.775e+02 |
| 5.000e-03 | -1.060e-04 | 6.470e-04 | -3.471e+03 | -2.271e+02 | 5.643e+02 |
| 1.000e-02 | -9.800e-05 | 9.390e-04 | -1.915e+03 | -9.587e+01 | 2.653e+02 |
| 1.500e-02 | -2.480e-04 | 2.440e-03 | -1.194e+03 | -8.038e+01 | 2.012e+02 |
| 2.000e-02 | -8.330e-04 | 1.938e-03 | -9.970e+02 | -3.493e+01 | 1.321e+02 |

(a) Sampling scheme A: 20 chromosomes drawn at 60 sampling time points.

| Selection coefficient | | | Allele age | | |
|---|---|---|---|---|---|
| True value | Bias | RMSE | True value | Bias | RMSE |
| 0.000e-00 | 4.540e-04 | 1.174e-03 | -6.999e+03 | 1.670e+03 | 6.169e+03 |
| 2.500e-03 | -4.000e-06 | 5.290e-04 | -5.951e+03 | -4.422e+02 | 9.792e+02 |
| 5.000e-03 | -4.900e-05 | 6.360e-04 | -3.471e+03 | -2.113e+02 | 5.677e+02 |
| 1.000e-02 | -2.920e-04 | 9.420e-04 | -1.915e+03 | -1.107e+02 | 2.641e+02 |
| 1.500e-02 | -2.750e-04 | 2.421e-03 | -1.194e+03 | -7.455e+01 | 1.867e+02 |
| 2.000e-02 | -7.320e-04 | 1.667e-03 | -9.970e+02 | -3.246e+01 | 1.287e+02 |

(b) Sampling scheme B: 120 chromosomes drawn at 10 sampling time points.

Table S7: Bias and RMSE of estimates for 100 datasets simulated under a growth demographic history with population size $N(k) = 16000$ for $k \in \{k_0, k_0 + 1, \ldots, [k_0/2]\}$ and $N(k) = 32000$ otherwise, and dominance parameter $h = 0.5$. The estimates are obtained assuming the true demographic history.

| Selection coefficient | | | Allele age | | |
|---|---|---|---|---|---|
| True value | Bias | RMSE | True value | Bias | RMSE |
| 0.000e-00 | 1.900e-04 | 6.940e-04 | -6.999e+03 | 7.102e+03 | 1.360e+04 |
| 2.500e-03 | 1.900e-05 | 6.320e-04 | -5.951e+03 | 1.012e+02 | 9.720e+02 |
| 5.000e-03 | -1.140e-04 | 6.490e-04 | -3.471e+03 | 3.917e+01 | 5.537e+02 |
| 1.000e-02 | -1.200e-04 | 9.490e-04 | -1.915e+03 | 4.835e+01 | 2.565e+02 |
| 1.500e-02 | -2.770e-04 | 2.452e-03 | -1.194e+03 | 1.575e+01 | 1.971e+02 |
| 2.000e-02 | -8.120e-04 | 1.915e-03 | -9.970e+02 | 4.053e+01 | 1.416e+02 |

(a) Sampling scheme A: 20 chromosomes drawn at 60 sampling time points.

| Selection coefficient | | | Allele age | | |
|---|---|---|---|---|---|
| True value | Bias | RMSE | True value | Bias | RMSE |
| 0.000e-00 | 2.260e-04 | 7.870e-04 | -6.999e+03 | 5.952e+03 | 1.192e+04 |
| 2.500e-03 | -1.200e-05 | 5.080e-04 | -5.951e+03 | 7.123e+01 | 9.956e+02 |
| 5.000e-03 | -2.900e-05 | 6.220e-04 | -3.471e+03 | 6.784e+01 | 5.607e+02 |
| 1.000e-02 | -3.190e-04 | 9.590e-04 | -1.915e+03 | 3.082e+01 | 2.539e+02 |
| 1.500e-02 | -3.710e-04 | 2.442e-03 | -1.194e+03 | 1.850e+01 | 1.814e+02 |
| 2.000e-02 | -7.590e-04 | 1.669e-03 | -9.970e+02 | 4.251e+01 | 1.380e+02 |

(b) Sampling scheme B: 120 chromosomes drawn at 10 sampling time points.

Table S8: Bias and RMSE of estimates for 100 datasets simulated under a growth demographic history with population size $N(k) = 16000$ for $k \in \{k_0, k_0 + 1, \ldots, [k_0/2]\}$ and $N(k) = 32000$ otherwise, and dominance parameter $h = 0.5$. The estimates are obtained assuming a constant demographic history with the most recent population size $N(k) = 32000$ for $k \in \{k_0, k_0 + 1, \ldots, 0\}$.

| Selection coefficient | | | Allele age | | |
|---|---|---|---|---|---|
| True value | Bias | RMSE | True value | Bias | RMSE |
| 0.000e-00 | 4.870e-04 | 1.255e-03 | -6.999e+03 | 2.212e+03 | 6.564e+03 |
| 2.500e-03 | 5.400e-05 | 5.590e-04 | -5.951e+03 | -2.802e+02 | 9.817e+02 |
| 5.000e-03 | 3.450e-04 | 1.378e-03 | -3.471e+03 | -2.036e+02 | 5.826e+02 |
| 1.000e-02 | -1.990e-04 | 1.487e-03 | -1.915e+03 | -3.859e+01 | 2.659e+02 |
| 1.500e-02 | -6.070e-04 | 3.162e-03 | -1.194e+03 | -1.031e+02 | 2.093e+02 |
| 2.000e-02 | -5.990e-04 | 3.254e-03 | -9.970e+02 | -5.880e+01 | 2.154e+02 |

(a) Sampling scheme A: 20 chromosomes drawn at 60 sampling time points.

| Selection coefficient | | | Allele age | | |
|---|---|---|---|---|---|
| True value | Bias | RMSE | True value | Bias | RMSE |
| 0.000e-00 | 4.950e-04 | 1.185e-03 | -6.999e+03 | 2.163e+03 | 7.576e+03 |
| 2.500e-03 | 2.300e-05 | 6.170e-04 | -5.951e+03 | -3.041e+02 | 9.721e+02 |
| 5.000e-03 | 2.120e-04 | 1.179e-03 | -3.471e+03 | -2.315e+02 | 5.883e+02 |
| 1.000e-02 | -3.840e-04 | 1.350e-03 | -1.915e+03 | -6.227e+01 | 2.589e+02 |
| 1.500e-02 | -6.350e-04 | 3.013e-03 | -1.194e+03 | -1.008e+02 | 2.416e+02 |
| 2.000e-02 | -4.950e-04 | 2.916e-03 | -9.970e+02 | -6.914e+01 | 1.984e+02 |

(b) Sampling scheme B: 120 chromosomes drawn at 10 sampling time points.

Table S9: Bias and RMSE of estimates for 100 datasets simulated under a decline demographic history with population size $N(k) = 100000$ for $k \in \{k_0, k_0 + 1, \ldots, [k_0/2]\}$ and $N(k) = 16000$ otherwise, and dominance parameter $h = 0.5$. The estimates are obtained assuming the true demographic history.

| Selection coefficient | | | Allele age | | |
|---|---|---|---|---|---|
| True value | Bias | RMSE | True value | Bias | RMSE |
| 0.000e-00 | 1.378e-03 | 2.267e-03 | -6.999e+03 | -1.971e+03 | 2.138e+03 |
| 2.500e-03 | 8.400e-05 | 5.810e-04 | -5.951e+03 | -1.499e+03 | 1.603e+03 |
| 5.000e-03 | 4.180e-04 | 1.559e-03 | -3.471e+03 | -8.937e+02 | 9.582e+02 |
| 1.000e-02 | -1.300e-04 | 1.491e-03 | -1.915e+03 | -4.031e+02 | 4.560e+02 |
| 1.500e-02 | -5.460e-04 | 3.190e-03 | -1.194e+03 | -3.108e+02 | 3.385e+02 |
| 2.000e-02 | -5.310e-04 | 3.501e-03 | -9.970e+02 | -2.368e+02 | 2.703e+02 |

(a) Sampling scheme A: 20 chromosomes drawn at 60 sampling time points.

| Selection coefficient | | | Allele age | | |
|---|---|---|---|---|---|
| True value | Bias | RMSE | True value | Bias | RMSE |
| 0.000e-00 | 1.292e-03 | 2.099e-03 | -6.999e+03 | -1.844e+03 | 2.093e+03 |
| 2.500e-03 | 8.100e-05 | 6.630e-04 | -5.951e+03 | -1.442e+03 | 1.533e+03 |
| 5.000e-03 | 2.910e-04 | 1.387e-03 | -3.471e+03 | -8.624e+02 | 9.360e+02 |
| 1.000e-02 | -3.090e-04 | 1.390e-03 | -1.915e+03 | -4.139e+02 | 4.589e+02 |
| 1.500e-02 | -5.560e-04 | 3.135e-03 | -1.194e+03 | -3.152e+02 | 3.502e+02 |
| 2.000e-02 | -4.480e-04 | 2.914e-03 | -9.970e+02 | -2.344e+02 | 2.756e+02 |

(b) Sampling scheme B: 120 chromosomes drawn at 10 sampling time points.

Table S10: Bias and RMSE of estimates for 100 datasets simulated under a decline demographic history with population size $N(k) = 100000$ for $k \in \{k_0, k_0 + 1, \ldots, [k_0/2]\}$ and $N(k) = 16000$ otherwise, and dominance parameter $h = 0.5$. The estimates are obtained assuming a constant demographic history with the most recent population size $N(k) = 16000$ for $k \in \{k_0, k_0 + 1, \ldots, 0\}$.

| Selection coefficient | | | Dominance parameter | | | Allele age | | |
|---|---|---|---|---|---|---|---|---|
| True value | Bias | RMSE | True value | Bias | RMSE | True value | Bias | RMSE |
| 1.000e-02 | -2.590e-04 | 1.650e-03 | 0.000e-00 | -1.000e-02 | 7.071e-02 | -1.778e+03 | -2.281e+01 | 2.026e+02 |
| 1.000e-02 | -1.300e-05 | 1.379e-03 | 5.000e-01 | 2.500e-02 | 1.323e-01 | -1.915e+03 | -5.024e+01 | 3.514e+02 |
| 1.000e-02 | 1.783e-03 | 6.589e-03 | 1.000e-00 | 7.500e-02 | 2.291e-01 | -5.736e+03 | -2.960e+02 | 1.519e+03 |

(a) Sampling scheme A: 20 chromosomes drawn at 60 sampling time points.

| Selection coefficient | | | Dominance parameter | | | Allele age | | |
|---|---|---|---|---|---|---|---|---|
| True value | Bias | RMSE | True value | Bias | RMSE | True value | Bias | RMSE |
| 1.000e-02 | -4.230e-04 | 1.464e-03 | 0.000e-00 | -5.000e-03 | 5.000e-02 | -1.778e+03 | -4.562e+01 | 1.457e+02 |
| 1.000e-02 | -1.520e-04 | 1.306e-03 | 5.000e-01 | 5.000e-03 | 8.660e-02 | -1.915e+03 | -1.082e+02 | 3.705e+02 |
| 1.000e-02 | 4.120e-04 | 4.103e-03 | 1.000e-00 | 1.100e-01 | 2.646e-01 | -5.736e+03 | -6.653e+02 | 1.086e+03 |

(b) Sampling scheme B: 120 chromosomes drawn at 10 sampling time points.

Table S11: Bias and RMSE of estimates for 100 datasets simulated under a constant demographic history with population size $N(k) = 16000$ for $k \in \{k_0, k_0+1, \ldots, 0\}$, where the dominance parameter $h$ is estimated from one of three values $\{0, 0.5, 1\}$.

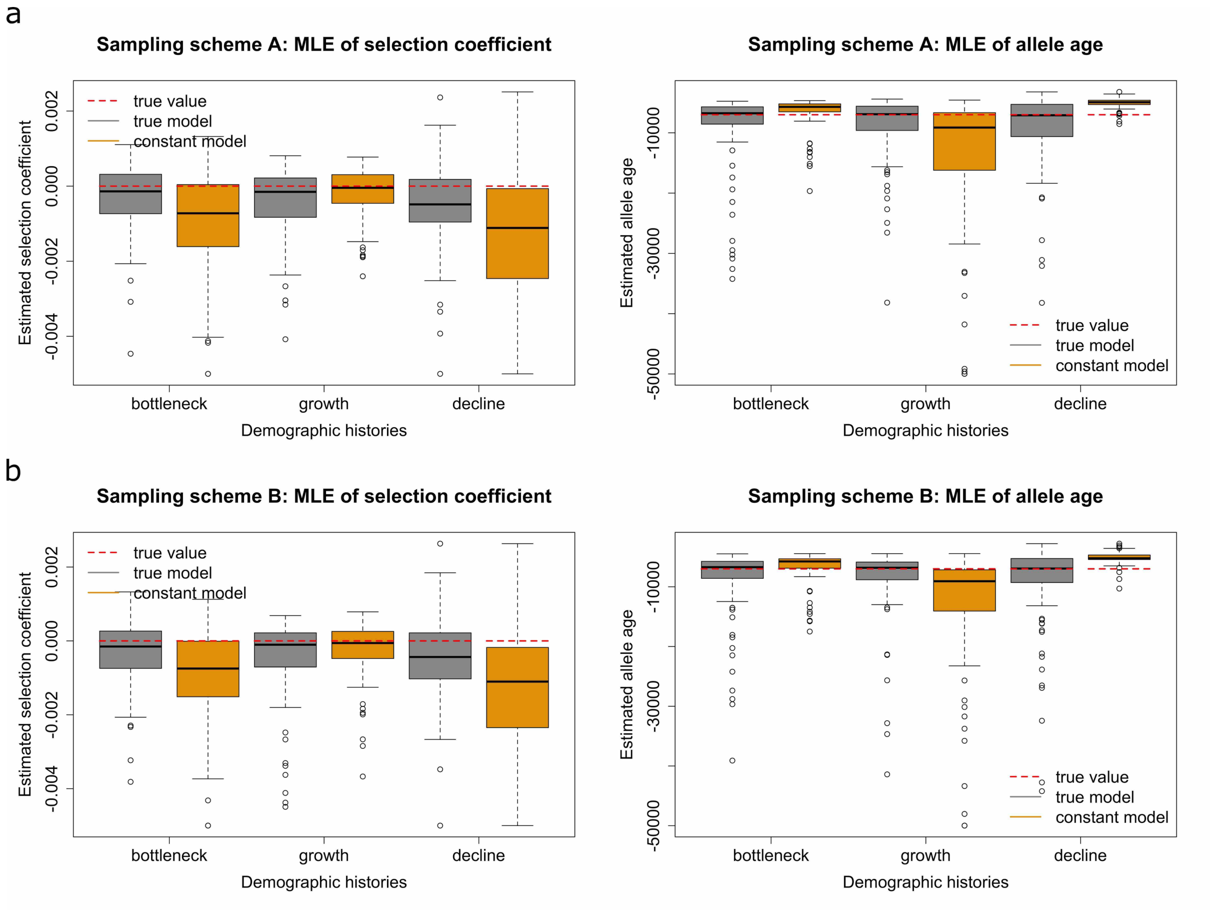


Figure S5: Empirical distributions of estimates for 100 datasets simulated under non-constant demographic histories with selection coefficient $s = 0$ (*i.e.*, *selectively neutrality*). Grey boxplots represent the estimates produced assuming the true demographic history, and yellow boxplots assuming a constant demographic history. Boxplots of the estimates for (a) sampling scheme A (b) sampling scheme B.

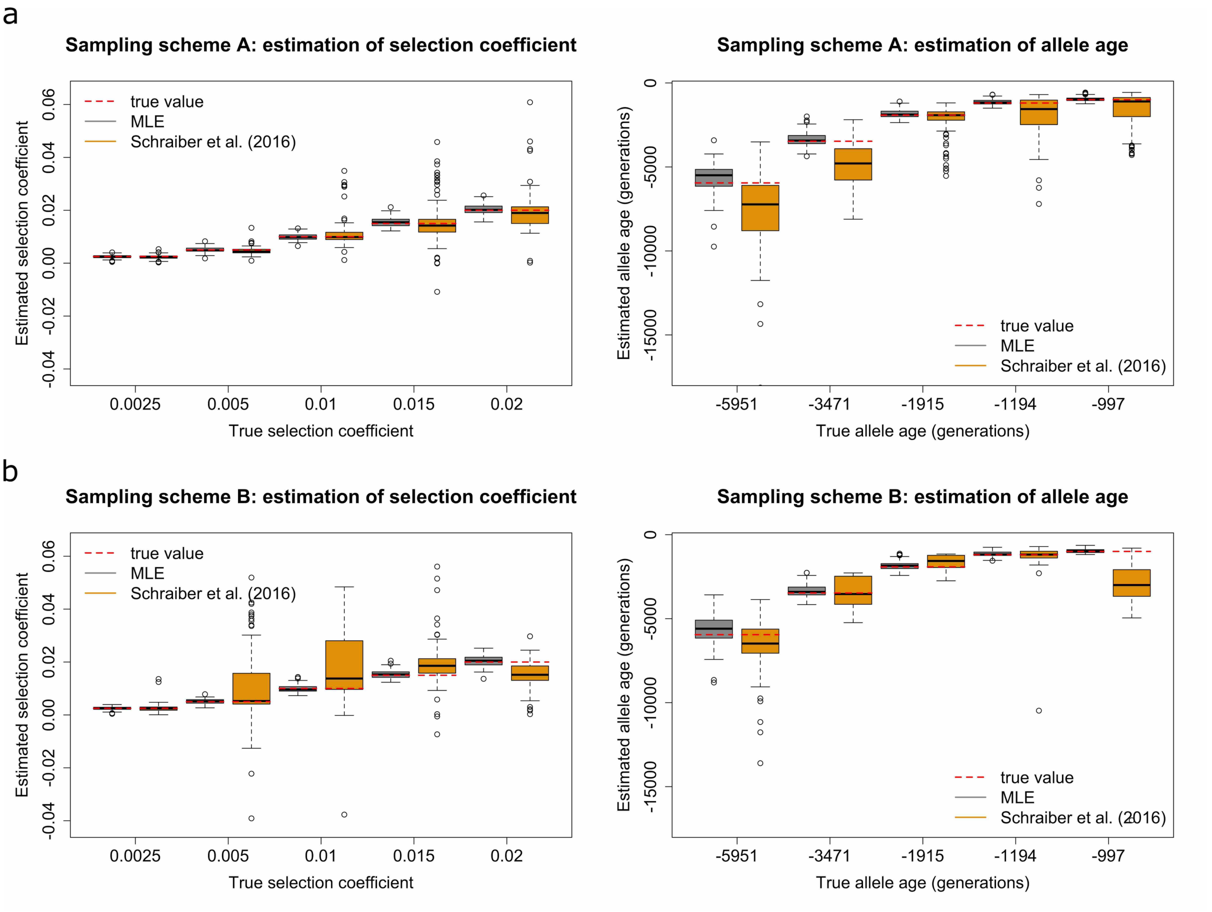


Figure S6: Empirical distributions of estimates for 100 datasets simulated under a bottleneck demographic history with population size $N(k) = 8000$ for $k \in \{[k_0/2], [k_0/2]+1, \ldots, [3k_0/4]\}$ and $N(k) = 16000$ otherwise, and dominance parameter $h = 0.5$. Grey boxplots represent the estimates produced with our method, and yellow boxplots using the approach of Schraiber et al. (2016). Boxplots of the estimates for (a) sampling scheme A (10 outliers of estimates for the selection coefficient and 1 outlier of estimates of the allele age using the method of Schraiber et al. (2016) lie outside the ranges of the plots and are not shown) (b) sampling scheme B (9 outliers of estimates for the selection coefficient and 1 outlier of estimates of the allele age using the method of Schraiber et al. (2016) lie outside the ranges of the plots and are not shown).

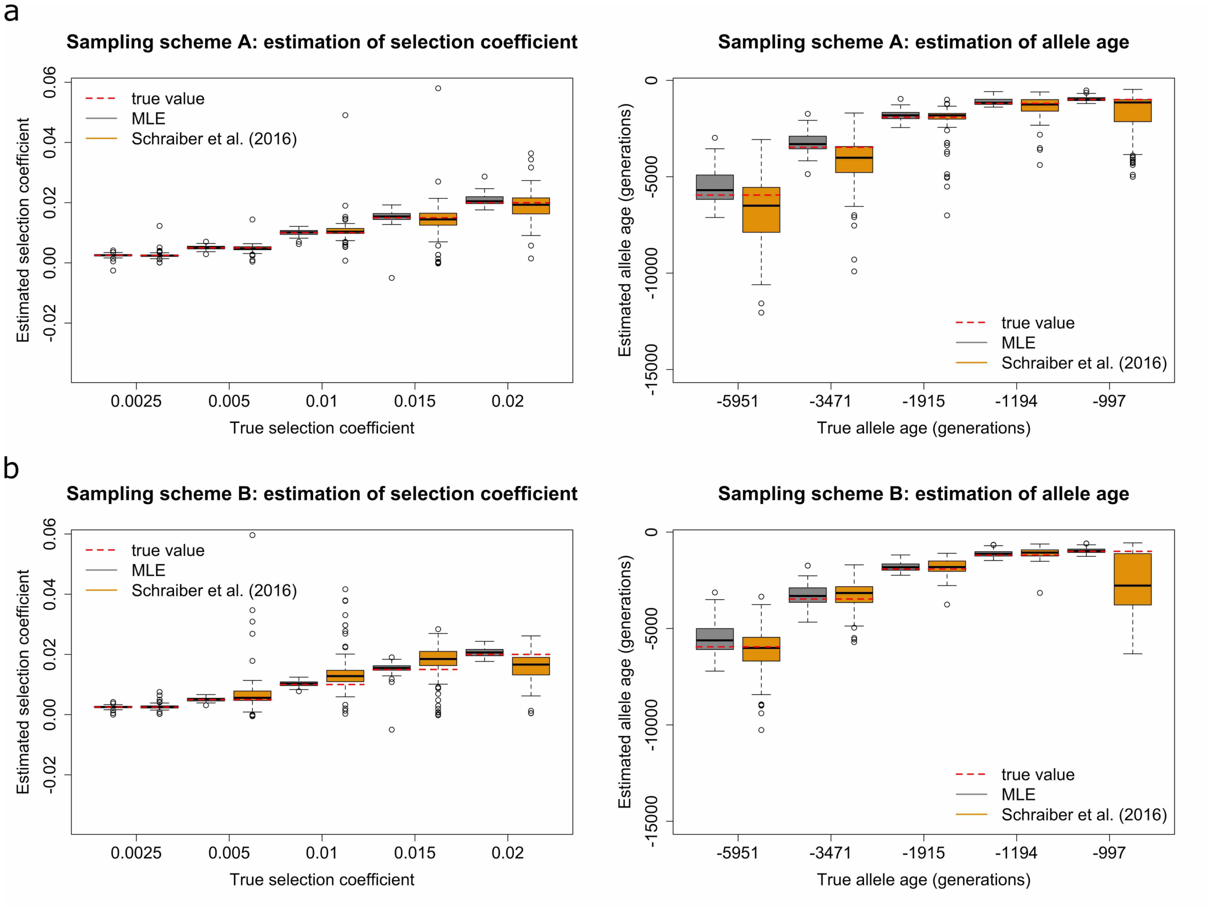


Figure S7: Empirical distributions of estimates for 100 datasets simulated under a growth demographic history with population size $N(k) = 16000$ for $k \in \{k_0, k_0 + 1, \dots, [k_0/2]\}$ and $N(k) = 32000$ otherwise, and dominance parameter $h = 0.5$. Grey boxplots represent the estimates produced with our method, and yellow boxplots using the approach of Schraiber et al. (2016). Boxplots of the estimates for (a) sampling scheme A (1 outlier of estimates for the selection coefficient using the method of Schraiber et al. (2016) lies outside the ranges of the plot and are not shown) (b) sampling scheme B (5 outliers of estimates for the selection coefficient using the method of Schraiber et al. (2016) lie outside the ranges of the plot and are not shown).

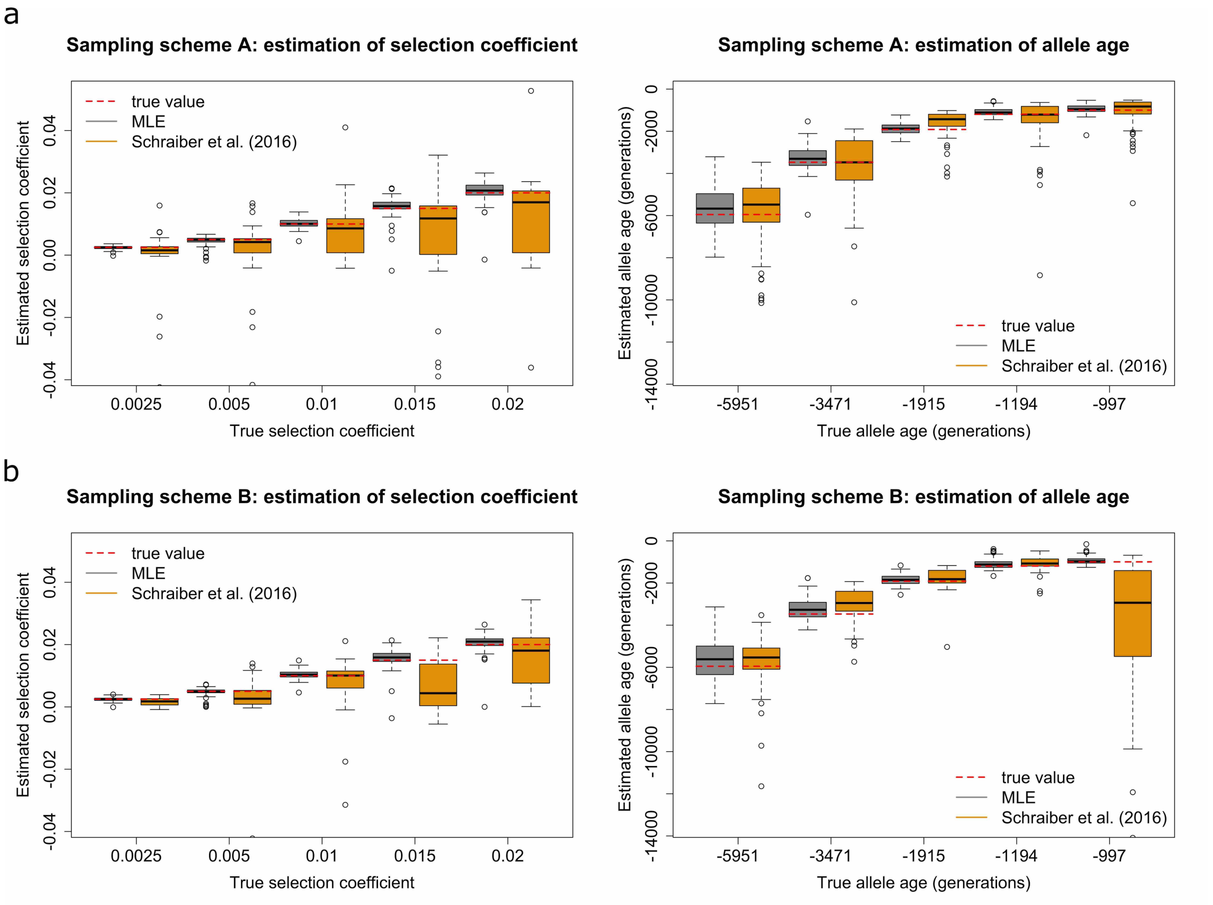


Figure S8: Empirical distributions of estimates for 100 datasets simulated under a decline demographic history with population size $N(k) = 100000$ for $k \in \{k_0, k_0 + 1, \ldots, [k_0/2]\}$ and $N(k) = 16000$ otherwise, and dominance parameter $h = 0.5$. Grey boxplots represent the estimates produced with our method, and yellow boxplots using the approach of Schraiber et al. (2016). Boxplots of the estimates for (a) sampling scheme A (89 outliers of estimates for the selection coefficient and 1 outlier of estimates of the allele age using the method of Schraiber et al. (2016) lie outside the ranges of the plots and are not shown) (b) sampling scheme B (12 outliers of estimates for the selection coefficient and 2 outliers of estimates of the allele age using the method of Schraiber et al. (2016) lie outside the ranges of the plots and are not shown).

| Selection coefficient | | | Allele age | | |
|---|---|---|---|---|---|
| True value | Bias | RMSE | True value | Bias | RMSE |
| 0.000e-00 | -2.260e-04 | 4.060e-04 | -6.999e+03 | 2.528e+03 | 3.655e+03 |
| 2.500e-03 | -8.100e-05 | 1.917e-03 | -5.951e+03 | 1.333e+03 | 2.574e+03 |
| 5.000e-03 | 1.280e-04 | 1.239e-03 | -3.471e+03 | 5.204e+02 | 1.381e+03 |
| 1.000e-02 | 6.670e-04 | 2.023e-03 | -1.915e+03 | 5.805e+02 | 1.142e+03 |
| 1.500e-02 | 9.370e-04 | 3.042e-03 | -1.277e+03 | 5.154e+02 | 9.012e+02 |
| 2.000e-02 | 2.197e-03 | 7.152e-03 | -9.970e+02 | 6.195e+02 | 1.056e+03 |

(a) Sampling scheme A: 20 chromosomes drawn at 60 sampling time points.

| Selection coefficient | | | Allele age | | |
|---|---|---|---|---|---|
| True value | Bias | RMSE | True value | Bias | RMSE |
| 0.000e-00 | -1.720e-04 | 2.040e-04 | -6.999e+03 | 1.683e+03 | 2.392e+03 |
| 2.500e-03 | -1.180e-04 | 2.064e-03 | -5.951e+03 | 5.990e+02 | 1.552e+03 |
| 5.000e-03 | -2.670e-04 | 1.789e-03 | -3.471e+03 | 6.255e+02 | 1.138e+03 |
| 1.000e-02 | 4.320e-04 | 1.909e-03 | -1.915e+03 | 9.491e+02 | 1.361e+03 |
| 1.500e-02 | 1.198e-03 | 5.593e-03 | -1.277e+03 | 1.475e+03 | 2.677e+03 |
| 2.000e-02 | 7.670e-04 | 4.403e-03 | -9.970e+02 | 1.815e+03 | 2.636e+03 |

(b) Sampling scheme B: 120 chromosomes drawn at 10 sampling time points.

Table S12: Bias and RMSE of estimates for 100 datasets simulated under a constant demographic history with population size $N(k) = 16000$ for $k \in \{k_0, k_0 + 1, \ldots, 0\}$ and dominance parameter $h = 0.5$. The estimates are obtained using the approach of Schraiber et al. (2016).

| Selection coefficient | | | Allele age | | |
|---|---|---|---|---|---|
| True value | Bias | RMSE | True value | Bias | RMSE |
| 0.000e-00 | -1.550e-04 | 1.940e-04 | -6.999e+03 | 2.847e+03 | 3.672e+03 |
| 2.500e-03 | 1.980e-04 | 8.550e-04 | -5.951e+03 | 1.633e+03 | 2.746e+03 |
| 5.000e-03 | 3.440e-04 | 1.439e-03 | -3.471e+03 | 1.436e+03 | 1.934e+03 |
| 1.000e-02 | 2.607e-03 | 2.800e-02 | -1.915e+03 | 2.889e+02 | 9.795e+02 |
| 1.500e-02 | 4.674e-02 | 3.946e-01 | -1.194e+03 | 7.202e+02 | 1.427e+03 |
| 2.000e-02 | 5.518e-02 | 5.559e-01 | -9.970e+02 | 5.756e+02 | 1.214e+03 |

(a) Sampling scheme A: 20 chromosomes drawn at 60 sampling time points.

| Selection coefficient | | | Allele age | | |
|---|---|---|---|---|---|
| True value | Bias | RMSE | True value | Bias | RMSE |
| 0.000e-00 | -1.780e-04 | 2.410e-04 | -6.999e+03 | 1.887e+03 | 2.497e+03 |
| 2.500e-03 | -4.700e-05 | 1.750e-03 | -5.951e+03 | 6.294e+02 | 1.605e+03 |
| 5.000e-03 | -3.559e-03 | 4.625e-02 | -3.471e+03 | 3.375e+00 | 8.837e+02 |
| 1.000e-02 | -1.496e-02 | 2.957e-01 | -1.915e+03 | -2.769e+02 | 5.168e+02 |
| 1.500e-02 | -3.780e-02 | 2.442e-01 | -1.194e+03 | 8.300e+01 | 9.729e+02 |
| 2.000e-02 | 4.597e-03 | 6.549e-03 | -9.970e+02 | 2.298e+03 | 4.136e+03 |

(b) Sampling scheme B: 120 chromosomes drawn at 10 sampling time points.

Table S13: Bias and RMSE of estimates for 100 datasets simulated under a bottleneck demographic history with population size $N(k) = 8000$ for $k \in \{[k_0/2], [k_0/2]+1, \ldots, [3k_0/4]\}$ and $N(k) = 16000$ otherwise, and dominance parameter $h = 0.5$. The estimates are obtained using the approach of Schraiber et al. (2016).

| Selection coefficient | | | Allele age | | |
|---|---|---|---|---|---|
| True value | Bias | RMSE | True value | Bias | RMSE |
| 0.000e-00 | -1.540e-04 | 2.470e-04 | -6.999e+03 | 2.945e+03 | 4.109e+03 |
| 2.500e-03 | -5.000e-06 | 1.210e-03 | -5.951e+03 | 8.148e+02 | 1.982e+03 |
| 5.000e-03 | 2.460e-04 | 1.385e-03 | -3.471e+03 | 8.109e+02 | 1.633e+03 |
| 1.000e-02 | -7.270e-04 | 4.488e-03 | -1.915e+03 | 1.755e+02 | 9.229e+02 |
| 1.500e-02 | 1.164e-03 | 7.155e-03 | -1.194e+03 | 1.791e+02 | 6.459e+02 |
| 2.000e-02 | 1.071e-03 | 5.079e-03 | -9.970e+02 | 7.165e+02 | 1.423e+03 |

(a) Sampling scheme A: 20 chromosomes drawn at 60 sampling time points.

| Selection coefficient | | | Allele age | | |
|---|---|---|---|---|---|
| True value | Bias | RMSE | True value | Bias | RMSE |
| 0.000e-00 | -1.460e-04 | 1.840e-04 | -6.999e+03 | 1.974e+03 | 2.613e+03 |
| 2.500e-03 | -9.200e-05 | 9.390e-04 | -5.951e+03 | 1.888e+02 | 1.174e+03 |
| 5.000e-03 | -5.245e-03 | 1.960e-02 | -3.471e+03 | -1.564e+02 | 7.836e+02 |
| 1.000e-02 | 1.780e-03 | 6.020e-02 | -1.915e+03 | -9.912e+01 | 4.161e+02 |
| 1.500e-02 | -2.571e-03 | 6.391e-03 | -1.194e+03 | -9.143e+01 | 2.911e+02 |
| 2.000e-02 | 4.373e-03 | 6.751e-03 | -9.970e+02 | 1.621e+03 | 2.185e+03 |

(b) Sampling scheme B: 120 chromosomes drawn at 10 sampling time points.

Table S14: Bias and RMSE of estimates for 100 datasets simulated under a growth demographic history with population size $N(k) = 16000$ for $k \in \{k_0, k_0 + 1, \ldots, [k_0/2]\}$ and $N(k) = 32000$ otherwise, and dominance parameter $h = 0.5$. The estimates are obtained using the approach of Schraiber et al. (2016).

| Selection coefficient | | | Allele age | | |
|---|---|---|---|---|---|
| True value | Bias | RMSE | True value | Bias | RMSE |
| 0.000e-00 | -8.110e-04 | 2.662e-02 | -6.999e+03 | 1.828e+03 | 3.646e+03 |
| 2.500e-03 | 8.853e-03 | 4.590e-02 | -5.951e+03 | -1.796e+02 | 1.575e+03 |
| 5.000e-03 | 3.623e-01 | 2.986e-00 | -3.471e+03 | 1.960e+02 | 1.419e+03 |
| 1.000e-02 | -3.927e-01 | 2.533e-00 | -1.915e+03 | -3.278e+02 | 6.558e+02 |
| 1.500e-02 | -1.776e-01 | 1.657e-00 | -1.194e+03 | 3.793e+02 | 1.800e+03 |
| 2.000e-02 | -1.263e-01 | 1.768e-00 | -9.970e+02 | 5.257e+01 | 6.919e+02 |

(a) Sampling scheme A: 20 chromosomes drawn at 60 sampling time points.

| Selection coefficient | | | Allele age | | |
|---|---|---|---|---|---|
| True value | Bias | RMSE | True value | Bias | RMSE |
| 0.000e-00 | -1.880e-04 | 4.240e-04 | -6.999e+03 | 5.317e+02 | 2.068e+03 |
| 2.500e-03 | 8.030e-04 | 1.383e-03 | -5.951e+03 | -3.238e+02 | 1.163e+03 |
| 5.000e-03 | -2.412e-02 | 3.496e-01 | -3.471e+03 | -4.949e+02 | 8.252e+02 |
| 1.000e-02 | -3.168e-02 | 2.381e-01 | -1.915e+03 | -1.524e+02 | 4.901e+02 |
| 1.500e-02 | 4.936e-01 | 3.196e-00 | -1.194e+03 | -1.279e+02 | 3.300e+02 |
| 2.000e-02 | 4.356e-03 | 1.035e-02 | -9.970e+02 | 2.785e+03 | 4.102e+03 |

(b) Sampling scheme B: 120 chromosomes drawn at 10 sampling time points.

Table S15: Bias and RMSE of estimates for 100 datasets simulated under a decline demographic history with population size $N(k) = 100000$ for $k \in \{k_0, k_0 + 1, \ldots, [k_0/2]\}$ and $N(k) = 16000$ otherwise, and dominance parameter $h = 0.5$. The estimates are obtained using the approach of Schraiber et al. (2016).

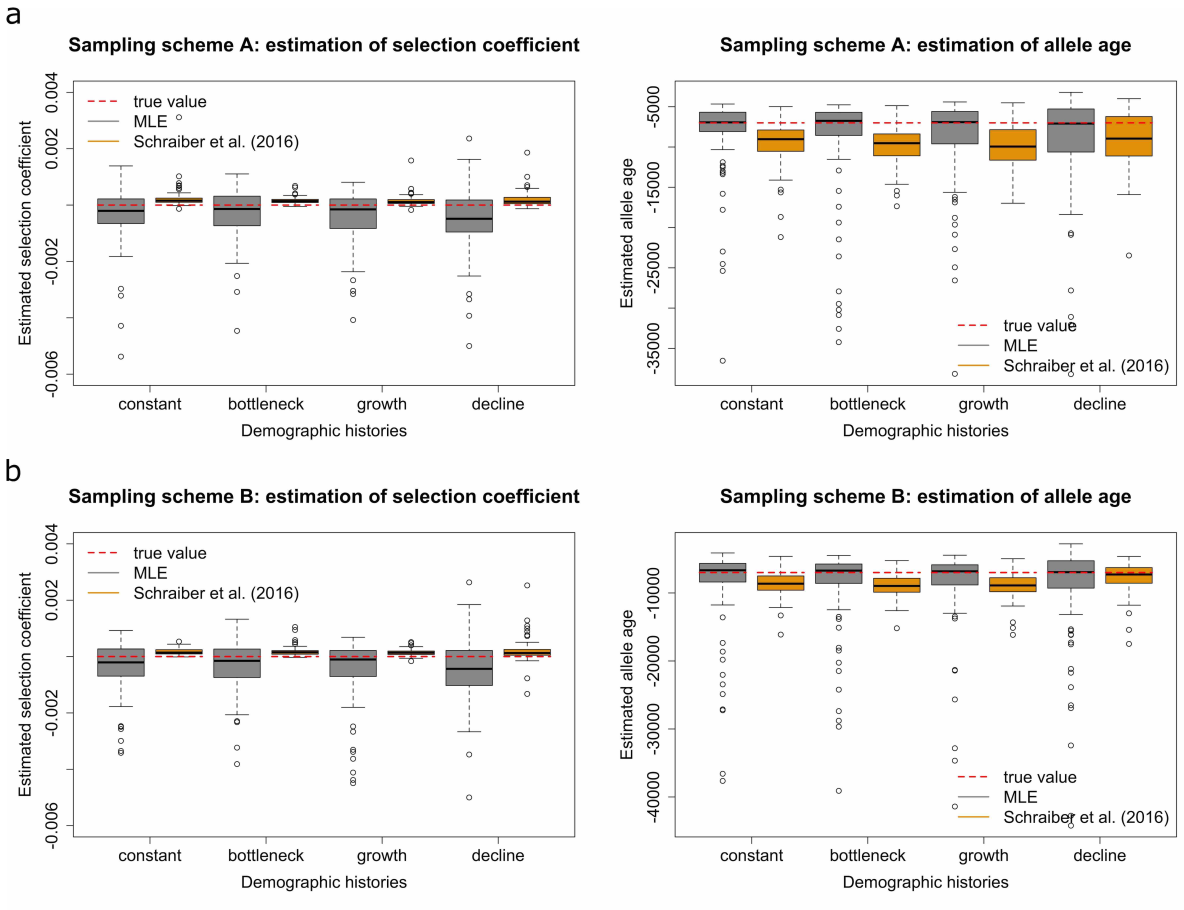


Figure S9: Empirical distributions of estimates for 100 datasets simulated under different demographic histories with selection coefficient $s = 0$ (*i.e.*, *selectively neutrality*). Grey boxplots represent the estimates produced with our method, and yellow boxplots using the approach of Schraiber et al. (2016). Boxplots of the estimates for (a) sampling scheme A (4 outliers of estimates for the selection coefficient, all under the decline demography, using the method of Schraiber et al. (2016) lie outside the ranges of the plot and are not shown) (b) sampling scheme B.

### File S3. Additional results on the analysis of real data

| Demographic history | Selection coefficient | Allele age |
|---|---|---|
| non-constant population size | 0.0013 [0.0004,0.0022] | -42982 [-175272,-18749] |
| constant population size of 16000 | 0.0018 [0.0008,0.0030] | -59224 [ -87974,-41008] |
| constant population size of 2500 | 0.0015 [0.0008,0.0022] | -16868 [ -25387,-16434] |

(a) *ASIP*

| Demographic history | Selection coefficient | Allele age |
|---|---|---|
| non-constant population size | 0.0126 [0.0091,0.0176] | -13644 [-15430,-12866] |
| constant population size of 16000 | 0.0129 [0.0088,0.0187] | -19942 [-26692,-16498] |
| constant population size of 2500 | 0.0115 [0.0077,0.0161] | -8716 [-11508, -7002] |

(b) *MC1R*

Table S16: Maximum likelihood estimates, as well as the 95% confidence intervals, for *ASIP* and *MC1R* under different demographic histories. The non-constant demographic history is given by Der Sarkissian et al. (2015).
